# 3D micropatterned traction force microscopy: a technique to control three-dimensional cell shape while measuring cell-substrate force transmission

**DOI:** 10.1101/2024.07.10.602889

**Authors:** Laura M. Faure, Manuel Gómez-González, Ona Baguer, Jordi Comelles, Elena Martínez, Marino Arroyo, Xavier Trepat, Pere Roca-Cusachs

**Author notes:** Equal authorship (Laura M. Faure, Manuel Gómez-González).

## Abstract

Cell shape and function are intimately linked, in a way that is mediated by the forces exerted between cells and their environment. The relationship between cell shape and forces has been extensively studied for cells seeded on flat 2-dimensional (2D) substrates, but not for cells in more physiological three-dimensional (3D) settings. Here, we demonstrate a technique called 3D micropatterned traction force microscopy (3D-μTFM) to confine cells in three-dimensional wells of defined shape, while simultaneously measuring the forces transmitted between cells and their microenvironment. This technique is based on the 3D micropatterning of polyacrylamide wells and on the calculation of 3D traction force from their deformation. With 3D-μTFM, we show that MCF10A breast epithelial cells exert defined, reproducible patterns of forces on their microenvironment, which can be both contractile and extensile. We further show that cells switch from a global contractile to extensile behaviour as their volume is reduced. Our technique enables the quantitative study of cell mechanobiology with full access to 3D cellular forces while having accurate control over cell morphology and the mechanical conditions of the microenvironment.

## Introduction

Cells within tissues exert forces on each other and on their microenvironment, modifying their shape and that of the tissue. This mechanical control of cell shape regulates developmental processes from gastrulation to organogenesis, disease progression, and aging^[1–3]^. Thus, the relationship between cell shape and cell mechanical forces is of major importance and has been extensively studied on two-dimensional (2D) surfaces. By combining cell micropatterning with force-measuring techniques such as traction force microscopy (TFM) or micropillars, researchers established a correlation between cell spreading area and the forces cells generate^[4–7]^. However, cells spreading on 2D surfaces do not adequately model the shape found in several physiological conditions highlighting the need for a system to control cell shape in three dimensions (3D) while measuring cell mechanical forces.

TFM, based on measuring the displacement of fiducial markers embedded in an elastic substrate, is one of the main techniques used to measure how cells exert forces on their surroundings. This technique was originally developed to measure 2D forces generated by single cells on flat surfaces but has since been extended to measure 3D forces produced by single cells^[8,9]^, cell monolayers^[10]^, and single cells or groups of cells embedded inside hydrogels in 3D^[11–13]^. However, none of the existing systems allow for the 3D control of cell shape while measuring cell forces, even though this relationship is fundamental in *vivo*^10^. In this work, we present a technique to compute traction forces generated by single cells in 3D micro-structures, 3D micropatterned traction force microscopy (3D-μTFM), and we apply it to study the relationship between cell shape and force in 3D. We show that epithelial cells display specific force patterns depending on their volume. While larger cells exert mostly contractile forces consistently with what has been described on 2D substrates, smaller cells generate extensile forces on their microenvironment through their actin cytoskeleton.

## Results

### Measuring 3D forces while controlling cell morphology

We used microfabrication techniques^[14,15]^ to develop a structured hydrogel to control cell morphology while enabling the measurement of 3D traction forces. As the material for our hydrogels, we chose polyacrylamide due to its linear elastic behaviour, transparency, tuneability, and widespread use in TFM on flat surfaces^[16–18]^ (Figure 1A). By polymerizing gels in contact with a poly(dimethylsiloxane) (PDMS) mold containing pillars, we generated wells in our gels with a size comparable to that of single cells. To tune cell size, we used pillars of 9 μm in height and either 15 μm or 19 μm in diameter. Because of gel swelling after polymerization, the resulting well dimensions were slightly different than the molds: 11-12 μm in height, and 10-11 μm or 14-16 μm in diameter, respectively. We refer to them as small and large wells, with volumes of 1060 μm^3^ ± 10 % and 1930 μm^3^ ± 9 % (mean ± standard deviation). Smaller and larger wells than these were also prepared, but our cellular model (MCF10A breast epithelial cells) either did not fit in very small wells or did not completely fill very large wells. Thus, these wells were discarded in further experiments. Fluorescent microbeads were added to the gel mixture as fiducial markers for measuring gel deformations. Beads accumulated near the free surfaces of the gel during polymerization. This accumulation was beneficial, as the distribution of beads on a thin layer enhances the accuracy of surface displacement measurements compared to a uniform, bulk bead distribution. Finally, we provided cell ligands to the wells by covalent binding of fibronectin to the gel surface.

**Figure 1:**
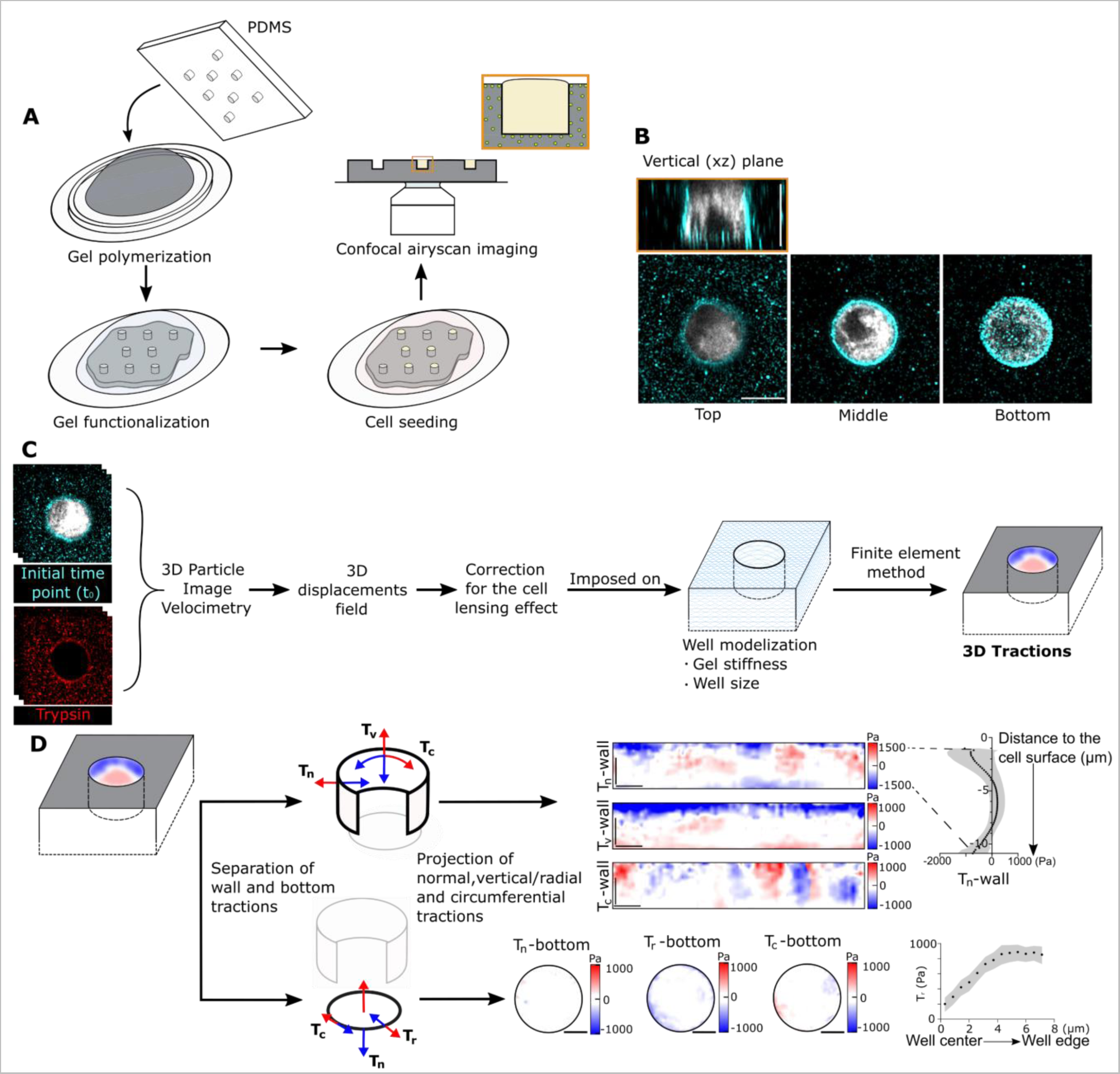
Workflow to measure 3D forces generated by cells of controlled morphologies. (A) Production of the experimental set-up. (B) Example of a cell (in white, labeled with CellTracker^TM)^ in a well, fluorescent beads are shown in cyan. Scale bar: 10 µm. (C) Pipeline analysis. For better visualization of the bead displacements generated by cells, the images of the beads in the initial and relaxed positions (before/after trypsinization) are artificially displayed in cyan and red, respectively. (D) From 3D tractions in the wells, maps of normal, vertical, and circumferential tractions to the wall (T_n_-wall, T_v_-wall, and T_c_-wall, respectively) to the wall were calculated, and maps of normal, radial, and circumferential tractions to the bottom of the well (T_n_-bottom, T_r_-bottom, and T_c_-bottom, respectively) were computed. From these maps, mean profiles of tractions were obtained and represented along the z-axis for the wall of the well or radially for the bottom of the well. See arrow colors for directions of positive/negative tractions in each case. Scale bar: 10 µm.

Having obtained control over cell shape, we proceeded to image our system to compute 3D traction forces. To achieve an optimal compromise between spatial and temporal resolution, we used a fast airy-scan confocal microscope (Carl Zeiss Ltd.). Following a workflow analogous to 2D-TFM (Figure 1C) we first set up a particle image velocimetry (3D PIV) algorithm to measure the 3D displacements of the fluorescent beads. Our 3D PIV compares a 3D stack image of the deformed gel with a 3D stack of the relaxed state, providing a 3D deformation map. Only displacements of the surface of the gel, including the top surface, the well wall, and the well bottom, were needed for our calculation. Then, we developed an inverse method to compute the 3D tractions through finite element modelling (FEM), imposing the measured displacement on a geometrical model of our wells. For each well size, a model was created as a cylindrical cavity of the same radius and depth as the imaged well. The model was meshed, and the mesh nodes of the upper surface were aligned with the 3D stack image of the gel relaxed state. The 3D tractions at the gel-cell interface were then obtained by solving an inverse FEM problem minimizing the discrepancy between computational and measured 3D displacements, introducing mechanical equilibrium as constraints in the optimization (Supplementary Information - note 1).

To represent the computed tractions, we took advantage of the cylindrical symmetry of the wells. We unfolded the traction fields exerted on the lateral wall and represented them as a rectangle, whereas we represented the traction fields exerted on the well bottom as a disk (Figure 1D). For better visualization and comparison of traction results, we decomposed the total tractions in normal, vertical, and circumferential traction maps for the wall (T_n_-wall, T_v_-wall and T_c_-wall, respectively) and normal, radial, and circumferential traction maps for the bottom (T_n_-bottom, T_r_-bottom, and T_c_-bottom, respectively) (Figure 1D). From these maps, we average the traction components along the well circumference and represent those averages as a function of the radial distance from the centre of the well (for the bottom surface) and as a function of the vertical distance from the well top (for the lateral wall).

### Characterization of the system

To estimate noise levels and assess the effect of gel swelling during medium changes, we first carried out our experimental protocol using empty wells (Figure S1, Supporting Information). Wall tractions showed values generally below 100 Pa, and no distinct pattern of forces on the maps except for peaks of 250 Pa at the top edge of T_n_-wall. In bottom tractions, peaks of 250Pa were also found at the edge. In both the walls and bottom of wells, edges are where we expected the largest discretization errors of the experimental well due to inaccuracies in precisely determining well shape (see methods). T_r_-bottom and T_c_-bottom maps showed dipole patterns with values around 500 Pa, associated with residual misalignment of the images (Figure S1Q-X, Supporting Information). Except for the edge effects in wall and bottom, all traction noises largely canceled out when computing average profiles along the wall height, or the bottom radius (Figure S1, Supporting Information). Comparing these average noise profiles to those of real cells (described below), signal-to-noise ratios were above 9.

Another potential source of noise comes from cells themselves. Cells are known to induce lensing effects in microscopy images due to their shape, their nucleus, and to differences in refractive index between themselves (around 1.36) and their surrounding media (around 1.34)^[19]^. These effects, in the order of hundreds of nanometres, have been shown to create aberrations in the measurement of forces generated by cells placed on 2D micropillars^[19]^ and are generally more complex in 3D settings. One of the strengths of our method is that, because our wells have simple and reproducible geometries, we can theoretically estimate and experimentally measure the displacement introduced by the difference in refraction index. We were thus able to correct it before calculating 3D tractions through FEM (Supplementary information - note 2, Figure 1C, Figure S2-4 Supporting Information). After this correction, we measured traction forces in a condition where they should not be present, trypsinized cells (Figure S4, Supporting Information). As it considers all potential sources, we took this measurement as a more complete quantification of noise, and we use it for the rest of the manuscript. Using this quantification of noise, average traction patterns showed signal-to-noise levels above 3.5, except in specific conditions discussed below.

### Cells exert contractile forces when placed in a large PAA well

When cells are adhered on 2D surfaces^[16–18]^ or embedded in 3D gels^[11,12]^, they are well known to exert contractile forces. We thus tested if cells confined in cylindrical wells display similar mechanical behaviour. To do so, we seeded MCF10A in our large wells, and measured traction forces. Forces normal to the wall (T_n_-wall) showed a characteristic pattern that varies along the z-axis. From top to bottom, forces are directed inward, outward, and inward again with a higher magnitude for inward-directed forces (average maximum at 700 Pa), than for outward-directed forces (average maximum at 150 Pa, Figure 2A-C). Vertical forces along the wall (T_v_-wall) show downward-directed forces at the top of the wall (1000 Pa in average) and upward-directed forces at the bottom (300 Pa in average, Figure. 2H-I). Thus, cells contract the walls, pulling the wall top and bottom towards the centre. Radial bottom tractions (T_r_-bottom) showed a similar pattern of contractile forces, where cells pull the bottom edge towards the centre (700 Pa on average, Figure 2K-M).

**Figure 2:**
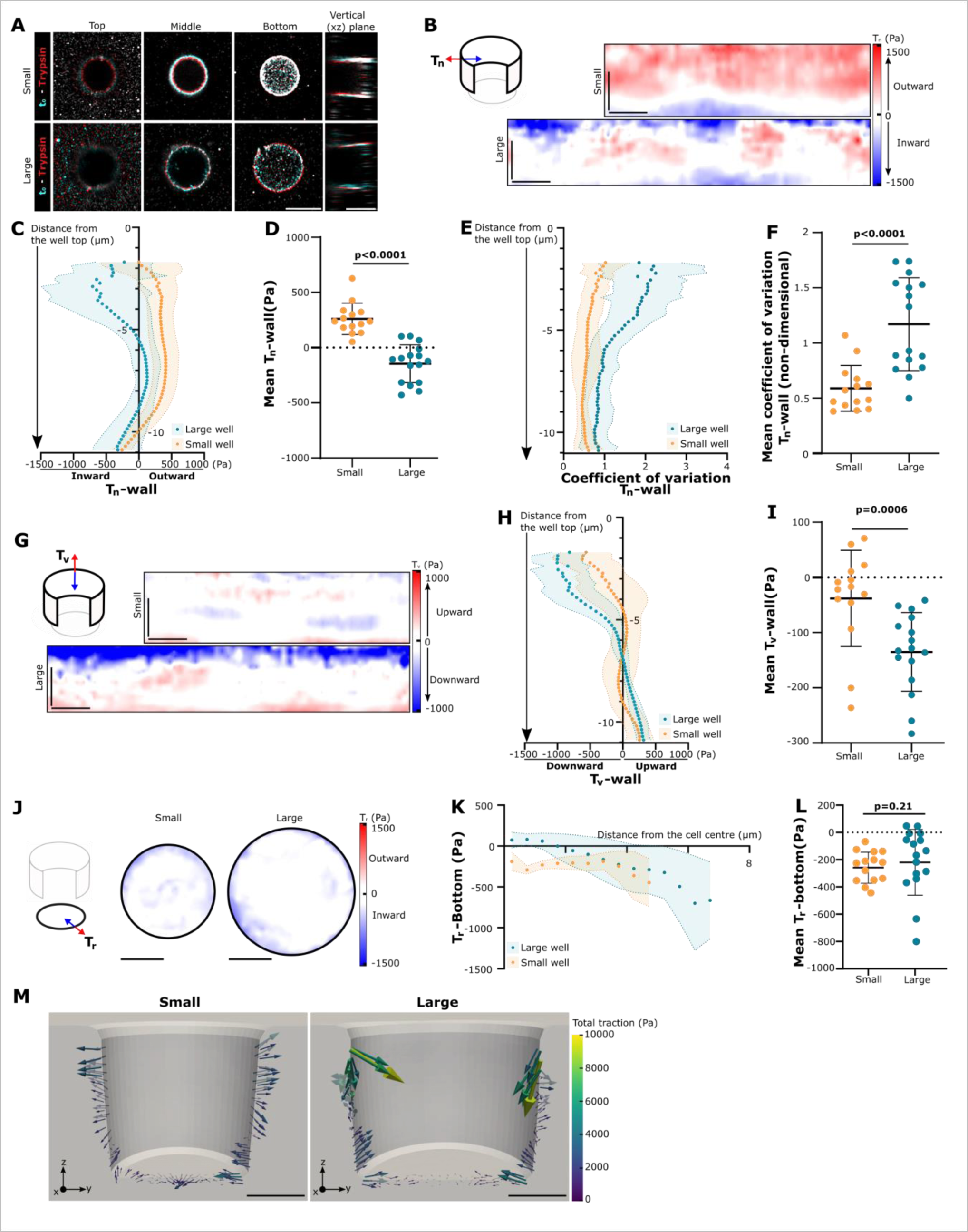
Cells push or pull depending on their volume. (A) Superposition of bead localization before (cyan) and after (red) trypsinization of example cells on small/large wells. Three xy planes of the stacks are shown (Top, Middle, and Bottom of the well), and a vertical plane (xz) of the same stacks. Scale bars: 10 µm. (B) Corresponding T_n_-wall maps. Scale bars: 5 µm. (C) Average T_n_-wall profiles along the z-axis. (D) T_n_-wall mean. p<0.0001, Mann-Whitney test. (E) Coefficient of variation of T_n_-wall profiles along the z-axis. (F) Mean of the coefficient of variation of T_n_-wall profiles. p<0.0001, unpaired T-test. (G) Example T_v_-wall maps. Scale bars: 5 µm. (H) Average T_v_-wall profiles along the z-axis. (I) T_v_-wall mean. p=0.0006, Mann-Whitney test. (J) Example T_r_-bottom maps. Scale bars: 5 µm. (K) Average radial profiles of the T_r_-bottom tractions. (L) T_r_-bottom mean. p=0.21, Mann-Whitney test. (M) Traction forces of a representative vertical cut from the small and large well data presented in panel a (Data displayed with Paraview). Scale bar: 5 µm. Data are mean ± standard deviation, n=14 and n=16 cells for the small and the large well conditions, respectively, from at least three independent experiments.

Normal and circumferential forces to the bottom of the well (T_n_-bottom and T_c_-bottom, respectively) are lower, with traction profiles below 100 Pa (Figure S5D-I, Supporting Information). Finally, circumferential forces along the wall (T_c_-wall) can reach 500 Pa but do not present a preferential directionality, as shown by mean profiles lower than 100 Pa. Average forces in these directions thus essentially measure noise and are only 30% higher than noise levels (Figure S5, Supporting Information). Moreover, their positioning does not correlate with any of the previous patterns from the T_n_-wall or the T_v_-wall. In conclusion, cells placed in the large wells present a characteristic traction pattern consisting of an apical and a basal ring of contractility which pull the well edges toward the cell centre (Figure 2M).

### Cell mechanical behaviour depends on cell volume

Next, we compared the large wells described above (volume of 1930 μm^3^ ± 9 %, mean ± standard deviation) to smaller wells (volume of 1060 μm^3^ ± 10 %, mean ± standard deviation). Forces normal to the wall (T_n_-wall) in cells seeded in small wells pointed outward along the top two-thirds of the wells (Figure 2A-C). Overall, the average of the T_n_-wall was positive for small wells but negative for large wells, meaning that cells in small wells exert extensile forces and not contractile forces (Figure 2D). Interestingly, T_n_-wall forces were also more homogeneous in small wells, as quantified by a lower coefficient of variation (standard deviation divided by the mean, Figure. 2E, F). This shows that cells in small wells lose the polar configuration with contractile peaks at the bottom and top, and rather exert a more homogeneous outward, extensile force.

Apart from T_n_-wall, other force patterns in small wells showed changes in their magnitude with respect to large wells, but not in their orientation. Cells placed in small wells still exert inward-directed forces at the bottom of the well but of slightly lower magnitude (500 Pa) than cells placed in large wells (700 Pa, Figure 2J-L). This is comparable to what was shown for cells placed on 2D patterns of different sizes^[4,5]^. Cells placed in the smaller wells also exert downward-directed forces at the top of the well and upward-directed forces at the bottom of the well, but of lower magnitude than for the large wells, with maximum values below 500 Pa (Figure 2H, I). As for large wells, T_c_-wall and T_c_-bottom maps display tractions generally below 400 Pa and 300 Pa, respectively, and no preferred directionality (Figure S5, Supporting Information). To conclude, confining cells to a smaller volume in the xy plane reduced overall force generation and changed cell mechanical behaviour from contractile to extensile (Figure 2M).

### Contractile behaviour in large wells depends on actin and myosin activity, while extensile behaviour in small wells requires actin polymerization

Cell generation of contractile forces has been widely studied and is mediated by the actomyosin cytoskeleton^[14,17,18]^. To assess the roles of both actin and myosin in the mechanical patterns observed in both small and large wells, we inhibited both actin polymerization with latrunculin A (LatA), and myosin contractility with blebbistatin (Bleb). To evaluate the effect of the drugs, we used our 3D-μTFM pipeline to compare the conditions before and after their addition (Figure 3A).

**Figure 3:**
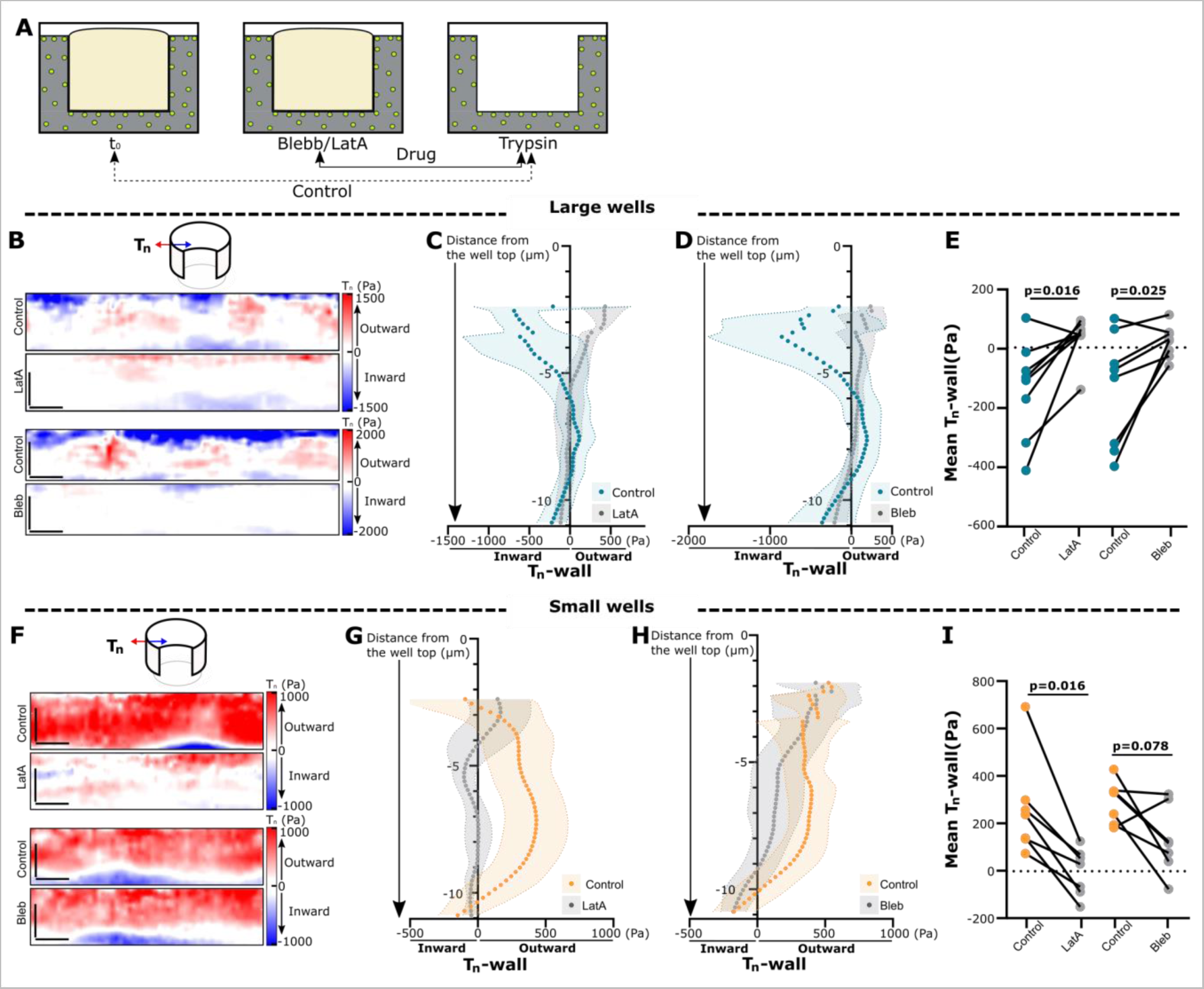
Contractile behaviour in large wells depends on actin and myosin activity, while extensile behaviour in small wells requires actin polymerization. (A) Schematic of the experimental set-up, bead positions in gels before any treatment (Control) or after the addition of blebbistatin (Bleb) or latrunculin A (LatA) are compared to bead positions after complete cell removal by trypsinization. (B, F) Example T_n_-wall maps comparing the blebbistatin/latrunculin A condition to the control for large (B), and small (F) wells. (C, D, G, H) Average T_n_-wall profiles along the z-axis for large (C, D), and small (G, H) wells. Data are mean ± standard deviation. (E, I) Example T_n_-wall means for large (E), and small (I) wells. Mean paired data are represented as connected. For large wells, Control/LatA: p=0.0016, Wilcoxon matched-pairs signed rank test; Control/Bleb: p=0.025, paired t-test. For small wells, Control/ LatA: p=0.016, Wilcoxon matched-pairs signed rank test; Control/Bleb: p=0.078, Wilcoxon matched-pairs signed rank test. n=8 for each large well condition, n=7 for each small well condition, from at least three independent experiments. Scale bars: 5 µm.a

We focus on T_n_-wall as it shows the highest forces and most relevant changes with cell size. In large wells, T_n_-wall maps are strongly affected by both latrunculin A and blebbistatin (Figure 3B). The maps and traction profiles in the latrunculin A and blebbistatin conditions show very low forces, with only residual traction at the top edge (Figure 3B-D) and a 70-75% reduction in tractions with respect to control conditions (Figure 3E). Besides, we observed similar trends in T_v_-wall and T_r_-bottom (Figure S6A-H, Supporting Information). In conclusion, and consistent with what has been shown on 2D surfaces, the contractile activity we observe in the large wells is due to the actomyosin cytoskeleton. Moreover, it is worth noticing that the inhibition of myosin contractility through the addition of blebbistatin is sufficient to flatten both inward-directed forces (contractile) and outward-directed forces (extensile) in the T_n_-wall (Figure 3D). Likely, the forces generated by the contractile top and bottom rings are strong enough to cause hydrostatic pressure pushing on the central part of the wall of the well.

In contrast to what we observe in large wells, the addition of latrunculin A or blebbistatin leads to different results in small wells. Latrunculin A reduced T_n_-wall forces almost completely (97% for the mean). However, blebbistatin induced a much milder trend towards force reduction in T_n_-wall forces (55% for the mean) which was not significant (Figure 3F-I). Similar trends are observed for T_r_-bottom (Figure S6M-P, Supporting Information) while neither blebbistatin nor latrunculin A induced a significant change on the T_v_-wall (Figure S6I-L, Supporting Information) which could be explained by the low force values already observed in case of the control. This indicates that actin polymerization but not myosin contractility generates most of the extensile forces exerted by cells placed in small wells.

## Discussion

In this study, we present a structured hydrogel system that allows for the measurement of forces generated by single cells of defined morphologies in 3D. 3D-TFM was demonstrated over 10 years ago^[8,9,11]^, and has been refined since^[12,13,20]^. However, previous approaches did not control cell shape, and one of their difficulties lies precisely in the proper detection of the cell contour, which heavily influences traction results and can create cell lensing effects. By using wells of defined shape, we not only removed this complexity in traction computation, but we also obtained very regular maps of traction, where the lensing effect was corrected. This allowed us to easily recognize and compare force patterns, such as the two contractile rings present in the large wells. Another issue in previous 3D-TFM approaches arose from the use of non-linear, degradable, or viscoelastic gels, which required important approximations to calculate forces^[12]^. We chose to prevent this by using polyacrylamide hydrogels, which are linear elastic, with known elastic properties. Finally, we note that confocal images do not present the same resolution in x/y and z, meaning that errors in displacements are larger along the z-axis. This generates asymmetry in the traction noise. By separating the total tractions into T_n_-wall, T_v_-wall, T_c_-wall, T_n_- bottom, T_r_-bottom, and T_c_-bottom we compare tractions that are more influenced by either the xy or z displacements, thus isolating the different components. Another solution would be to use light-sheet microscopes to image the set-ups, since it produces images with isotropic resolution, although lensing effects might be more severe.

The ability of cells to exert both pulling and pushing forces has been well described, although measuring pulling forces has proven to be easier. Pushing forces, exerted by extensile cellular structures such as lamellipodia, filopodia, and podosomes^[21–24]^, have typically been measured using low throughput techniques such as Atomic Force Microscopy (AFM), optical or magnetic tweezers, and glass fiber deflection. For example, AFM has been used to measure extensile forces by keratinocyte lamellipodia^[21]^ or by macrophage podosomes^[23,24]^ of the order of 10 nN. Recent findings on amoeboid migration have also highlighted the importance of protrusive forces at the cell leading edge, but related forces have not been measured, due to the complexity of the system, for cells fully embedded in 3D matrices^[25]^. In this study, we present a novel method that allowed us to uncover a new mechanical behaviour: an extensile force consistently applied over the cell diameter when cells are confined in a plane. We estimate that this force is around 90nN, which is significantly higher than what has been observed for pushing forces until now. This difference could be explained by the large area over which this pushing force is applied (500 µm^2^) when compared to lamellipodia (0.6 µm^2^ in^[21]^) or podosomes (1-12 µm^2^).

The ability of our system to measure extensile forces allowed us to determine that diminishing cell diameter from 14 µm to 11 µm is sufficient to switch cell mechanical behaviour from contractile to extensile. Forcing cells into smaller volumes may increase protein concentration, and this could affect actin organization. In turn, contractile forces exerted by myosin II motors depend on actin organization, specifically on the length of actin filaments^[26]^, the state of these filaments (bundled or branched^[27,28]^), and the ability of myosin II to enter the actin mesh^[29]^. Thus, we hypothesize that such effects on actomyosin organization could explain the observed mechanical effect of cell volume. Of note, this could explain the difference as well in the lensing effect detected between cells placed in small or large wells. Further studies are needed to verify this hypothesis.

Regardless of the mechanism, an overall switch from a contractile to an extensile behaviour may have relevant implications in tissues. In cell monolayers, cell-substrate forces have been systematically measured to be contractile^[30,31]^. Transmission of contractility can, in fact, span long distances across monolayers, enabling long-range mechanical force transmission, and downstream effects such as directed migration towards stiff tissues (durotaxis)^[31]^. In such systems, however, forces exerted by cells are only measurable on the substrate, and forces exerted in their mid-plane (where we see extensile forces in our setting) are inaccessible. Potentially, a switch from contractile to extensile behaviour induced by cell confinement would dampen overall contractile forces, preventing mechanical force transmission over long distances and ensuing mechanotransduction mechanisms.

How cells acquire and maintain their shape, how this impacts tissue shape, and the relationship between cell shape and function have been central questions in cell biology^[32–34]^. To answer them, researchers have developed different techniques. Microcontact printing enabled researchers to decipher the impact of cell spreading area in 2D on cell fate^[35]^, cellular organization^[36,37]^, or cell ability to generate forces^[4,5]^. Microniche systems were later developed to fully control cell shapes in 3D. Made of stiff substrates or structured hydrogels, these microniches can host from a single cell to an organoid and have been used to decipher shape/function relationships for example in hepatocyte doublets^[38]^ or stem cell colonies^[39]^. In recent years, they have become widely used, and have also become more refined ^[39,40]^. Our developed system is applicable to such microniche systems, either in single-cell settings as done here or in multicellular settings.

In conclusion, we developed a new system: 3D-μTFM, where we applied 3D-TFM to micro-structured gels to measure previously inaccessible force patterns, which depend on 3D cell morphology. This system can be combined with current microniche technologies, adding a mechanical dimension to the study of cell shape/function relationships in 3D.

## Methods

### Cell culture

Mammary epithelial cells (MCF10A) were purchased from ATCC. Cells were used for a maximum of 30 passages and were cultured in DMEM-F12 (LifeTechnologies, 21331-020) with 5% horse serum, 1% penicillin-streptomycin, EGF (20 ng/ml), hydrocortisone (0.5 μg/ml), cholera toxin (100 ng/ml), and insulin (10 μg/ml). All cells were regularly tested for mycoplasma contamination. When used, blebbistatin (Sigma Aldrich – B0560) was used at a concentration of 25 uM and Latrunculin A (Sigma Aldrich – L5163) at a concentration of 500 nM.

### Fabrication of microstructured polyacrylamide (PAA) hydrogels

Gels were prepared as described previously^[15]^. Briefly, PDMS prepolymer (Sylgard 184 Silicon Elastomer, Dow Corning) mixed at a ratio of 10:1 w/w was spun-coated onto a silicon mold containing wells of 15 or 19 µm of diameter spaced 100 µm from each other, forming thin membranes of 50 µm in thickness. Silicon molds were fabricated on silicon wafers made from SU8-50 using conventional photolithography. PDMS was then cured overnight at 65°C. After curing, PDMS membranes were cut into pieces including both structured and flat regions and put onto an 18 mm glass coverslip. 6 well – glass bottom MatTek were used to cast the hydrogels. First, glass silanization was performed to ensure a stable attachment of PAA hydrogels. To this end, glass coverslips were incubated for 15 min with 3-(Trimethoxysilyl)propyl methacrylate (Sigma Aldrich – 440159), acetic acid (Sigma Aldrich – 1612), and 96 % ethanol at a ratio of 1:1:14 v/v, rinsed three times with 96 % ethanol and dried.

PAA hydrogel mix was prepared as described previously^[17,18]^. For 15 kPa gels, 19 % v/v of acrylamide 40 % (Bio-rad - 161-0140) and 8 % v/v bis-acrylamide 2 % (Bio-rad: 1610142) were added to produce the gels and mixed with 2 % v/v fluorescent carboxylated 200 nm beads (Invitrogen), 0.5 % v/v APS (Sigma Aldrich - A3678), and 0.05 % v/v tetramethyl-ethylene-amine (Sigma Aldrich - T9281). To achieve covalent functionalization of the PAA hydrogels, we added acrylic acid (Sigma Aldrich - 147230) to the mix as a co-monomer of acrylamide and at a 2 % concentration of monomers. To cast the gels, a PDMS ring spacer (100-150um thick) was placed on the glass bottom of the MatTek dishes. 100 µl of gel mix was poured inside the rings and immediately covered by the 18 mm glass coverslip with the PDMS mold thus putting the mold in contact with the PAA hydrogel. PAA hydrogels were left to polymerize for 2 hours at room temperature. The polymerized hydrogels were de-molded by carefully removing the coverslip and then stored in PBS for later activation and functionalization.

For 150 kPa gel, 30 % v/v of acrylamide 40 % (Bio-rad - 161-0140) and 30 % v/v bis-acrylamide 2 % (Bio-rad: 1610142) were added to produce the gels and mixed with 2 % v/v fluorescent carboxylated 200 nm beads (Invitrogen) and 1 % v/v Irgacure 2959 (BASF). To achieve covalent functionalization of the PAA hydrogels, we added acrylic acid (Sigma - 147230), 0,15% v/v for 15 kPa gels and 0,45% v/v for 15 kPa gels. To cast the gels, 15 ul of gel mix was poured on the glass bottom of the MatTek dishes and covered with a 12 mm glass coverslip presenting the pillar patterns in Norland Optical Adhesive 81. PAA hydrogels were polymerized for 15 minutes under a UV lamp at room temperature. The polymerized hydrogels were de-molded by carefully removing the coverslip and then stored in PBS for later activation and functionalization.

### Functionalization of the structured PAA hydrogels

First, the acid groups present in the gel were activated through the EDC (1-Ethyl-3 (3’-dimethylaminopropyl) carbodiimide HCL, Merk - 8510070025) NHS (N-Hydroxysuccinimide, Sigma - 130672) chemistry. Briefly, 190 mg of EDC was mixed with 230 mg of NHS and diluted in 20mM HEPES, pH7 (Life Technologies: 15630056), added to the gel, and incubated at 37°C for 20 minutes. The gels were then rinsed once with PBS and incubated with a 30 µg/ml fibronectin solution (fibronectin from human plasma, F0895 Sigma Aldrich) for 1 hour at room temperature. Finally, the gels were rinsed and passivated using a poly(L-lysine) - G - poly(ethylene glycol) (Susos AG: PLL (20 kDa)-g[3,5]-PEG (2) solution at 0.1 mg/ml in PBS for 1 hour at room temperature.

### 3-dimensional Traction Force Microscopy - Imaging

Cells were seeded overnight on structured PAA hydrogels, fabricated as described above. The next morning, cells were labeled using red CellTracker (ThermoFisher Scientific – C34552) diluted at 0.03 % v/v in media for 25 minutes at 37°C.

Traction force experiments were carried out using the fast-Airyscan mode of an inverted Zeiss LSM880 confocal with a glycerin immersion 40X/1.2 objective equipped with control of temperature, CO_2_, and humidity. For each condition, small regions of interest were defined around wells that would present a cell. Only cells completely filling wells, and not spreading or protruding out of the wells, were selected for imaging and subsequent analysis. Additionally, a neighbouring region of interest containing an empty well was also analysed as a control of gel swelling. To obtain 3D images of the wells, stacks were taken with a z-step of 0.2 µm. Fluorescent images of the beads and cells were acquired at the initial time point, after the addition of the drug, and after cell trypsinization (only for the image of the beads).

### 3-dimensional Traction Force Microscopy - Analysis

All codes generated are available at https://github.com/xt-prc-lab/3D_Micropatterned_Traction_Force_Microscopy.git.

#### 3D-PIV

A confocal fluorescence microscopy stack was acquired spanning the whole depth of the well, both when deformed by a cell and after relaxation by trypsin or drugs, with a z-step of the objective (Δs) of 0.2 µm. The axial positional shift of the focal plane (Δf), due to the refraction index mismatch between the sample (n2) and the immersion medium of the objective (n1), was corrected via Visser’s formula ^[41]^

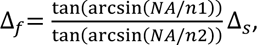

where NA is the numerical aperture of the objective. It is known that Visser’s formula might not be quantitatively accurate for some combinations of samples and objectives^[42]^, thus we checked its applicability by reconstructing the shape of spherical particles embedded in PAA gels.

The 3D displacement field of the whole gel around the well was calculated by implementing a custom 3D PIV in Matlab (MathWorks, Inc.)^[43,44]^. The whole stack was discretized in individual interrogation boxes, and their relative displacement between the deformed and reference configuration was calculated.

#### Traction inference using Finite Element Modelling

We first considered a direct approach, in which the measured 3D displacement field on the surface of the gel (top surface and well surface) was imposed as boundary conditions in a finite element model of the well. This approach was implemented in Matlab (MathWorks) and Abaqus (Dassault Systemes). Geometrically, we considered a square prism with the same height as the gel with a cylindrical cavity on the top surface of the same depth and average radius as the physical well. The edges of the cylinder were rounded using a radius of 0.5 μm for the bottom edge and of 1 μm for the top edge of the cylinder to closely follow the geometry of the physical wells. Each model was meshed in Abaqus with hybrid 4-node linear tetrahedron elements. We modelled the gel as a hyperelastic solid to account for possible geometrical and material nonlinearities. The geometry of the mesh was imported in Matlab, and the 3D displacement values of each node at the surface of the gel were interpolated from the 3D PIV displacement field. This approach, however, resulted in non-zero tractions at the top surface of the gel, although this part of the surface did not interact with cells. Imposing the 3D displacement only at the well surface fixes this issue but disregards experimental measurements at the top surface. To use all measured data while obtaining mechanically meaningful results consistent with force balance, we developed an inverse approach as detailed in Supplementary information - note 1.

#### Unfolding of the deformation and traction fields

The deformation field measured by the PIV was provided in a 3D volume around the well, without information on the location of the different surfaces of our system. To locate the upper surface of the gel, we calculated the mean intensity of each plane of the fluorescent bead microscopy image stack. The peak location of this distribution provided the plane of the gel’s surface. We did not find any consistent signature on the mean intensity signal pointing toward the location of the base of the well. If using the standard deviation of the intensity rather than the mean, the largest peak also indicated the location of the upper surface, while the second peak pointed towards the general location of the well’s bottom. However, this second peak contained too many errors in our measurements to be used. We thus located the bottom of the well with an offset, equal to the model’s depth, from the upper surface of the gel. To locate the surface of the vertical wall of the well, we first detected the well circumference in a central plane, either automatically or manually, and then its centre. The wall surface was then defined as a cylinder with the same radius as the well.

Given the regular shapes of the mesh of the model of the wells, the location of the different surfaces was readily available for the traction fields. The upper surface of the gel was composed of the nodes with the highest z-coordinate. The bottom surface of the well was composed of the nodes with the lowest z-coordinate, and the rest belonged to the wall of the well. It is important to notice that the nodes at the rounded area between the upper surface of the gel and the vertical wall of the well is where we had the highest discretization errors and geometrical uncertainties. With this in mind, we excluded the upper rounded area (around 1 μm) from the wall of the well region in the unfolded maps. The traction maps so defined are composed of discrete points (the nodes of the model). To provide a continuous representation, we interpolated the maps with Matlab’s 2D interpolation of scattered data, *scatteredInterpolant*, with the *natural neighbor interpolation* method.

### Statistical analysis and graphic representation

In the T_n_-wall, T_v_-wall and T_c_-wall profile representations, as some profiles were longer than others (12 µm deep wells compared to 11 µm deep wells) all profiles were aligned to the same lowest point. The same was done for the representation of the normal displacement to the wall of the well. Finally, when aligned, if only one profile was longer than the other, the extra values were not displayed as they were not associated with any standard deviation.

Statistical analyses were performed using GraphPad Prism software (version 8). Statistical significance was determined by the specific tests indicated in the corresponding figure legends. Non-parametric tests were performed when neither original nor log-10-transformed datasets were normally distributed. All the experiments presented here were repeated in at least three independent experiments.

## Acknowledgments

We thank A. Del campo for initial discussion, S. Usieto, A. Menéndez, N. Castro, M. Purciolas, and M Morcillo Sanchez for providing technical support; E. Latorre, I. Granero-Moya and M. Matejcic for providing data analysis tools; E. Dalaka for her AFM expertise, JB. Fiche, Z. Kechagia, A. Labernadie, A Beedle, A.L. Le Roux, L. Rosetti and R. Sunyer, as well as all the members of the groups of P.R.-C. and X.T. for helpful discussions.

## Author contributions

Conceptualization: L.M.F, P.R.-C.

Resources: L.M.F., M.G.G., J.C., E.M., X.T.

Data curation: L.M.F., M.G.G.

Software: M.G.G.

Formal analysis: L.M.F., M.G.G.

Supervision: M.A., X.T., P.R.-C.

Funding acquisition: E.M., M.A., X.T., P.R.-C.

Validation: L.M.F., M.G.G.

Investigation: L.M.F., M.G.G., O.B.C.

Visualization: L.M.F., M.G.G

Methodology: L.M.F., M.G.G., J.C.

Writing—original draft: L.M.F.

Project administration: X.T., P.R.-C.

Writing—review and editing: L.M.F., M.G.G., X.T., P.R.-C.

## Data availability

Source data shown in all figures is available as supplementary materials.

## Code availability

All generated codes are available at https://github.com/xt-prc-lab/3D_Micropatterned_Traction_Force_Microscopy.git.

