## Supplemental figures and notes for "3D micropatterned traction force microscopy: a technique to control three-dimensional cell shape while measuring cell-substrate force transmission"

**Inventory of Supplementary Information:**

- Supplementary Information – Figure S1: Quantification of traction noise in empty wells.
- Supplementary Information – Figure S2: Characterization of the “refractive” field of displacement induced by the cell lensing effect in 15 kPa gels.
- Supplementary Information – Figure S3: Characterization of the “refractive” field of displacement induced by the cell lensing effect in 150 kPa gels.
- Supplementary Information – Figure S4: Measurement of the traction noise in experiments corrected for the cell lensing effect.
- Supplementary Information – Figure S5: Wall circumferential tractions and bottom normal and circumferential tractions for cells placed on small or large wells.
- Supplementary Information – Figure S6: Wall vertical tractions and bottom radial tractions for cells placed on small or large wells.
- Supplementary Information – note 1: Finite element method for traction inference.
- Supplementary Information – note 2: Theoretical and experimental estimation of lensing effects and their correction.

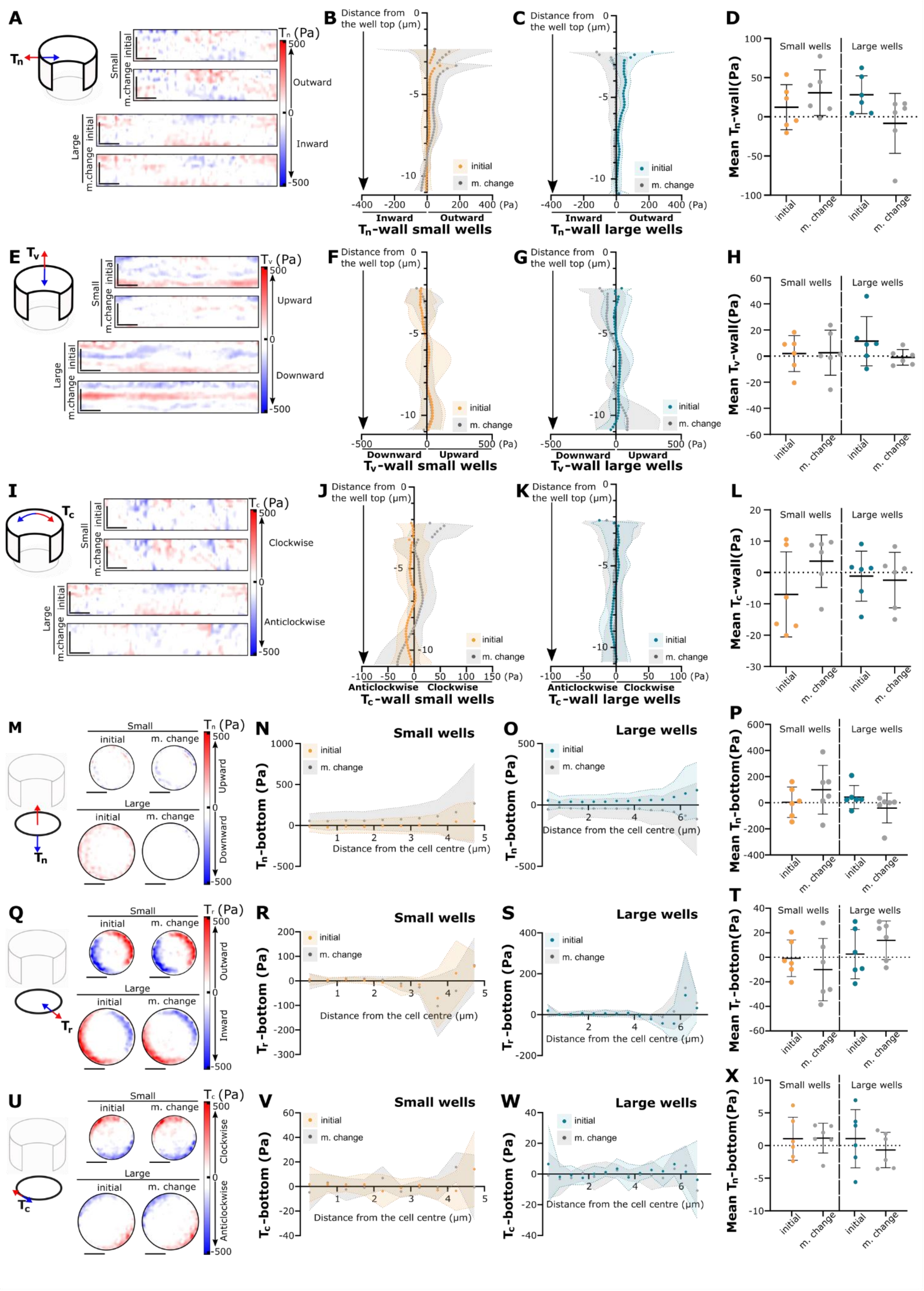


**Figure S1: Quantification of traction noise in empty wells.** (A) Examples of $T_n$-wall maps in empty wells, computed by comparing bead positions in the initial condition (initial) or after a first medium change (m. change, used to introduce drug treatments in experiments with cells), respectively, to the last medium change (used to trypsinize cells in experiments with cells). (B, C) Average $T_n$-wall profiles along the z-axis for the small (B) and large (C) wells. (D) $T_n$-wall means. (E) Examples of $T_v$-wall maps in empty wells. (F, G) Average $T_v$-wall profiles along the z-axis for the small (F) and large (G) wells. (H) $T_v$-wall means. (I) Examples of $T_c$-wall maps in empty wells. (J, K) Average $T_c$-wall profiles along the z-axis for the small (J) and large (K) wells. (L) $T_c$-wall means. (M) Examples of $T_n$-bottom maps in empty wells. (N, O) Average radial profiles of $T_n$-bottom for the small (N) and large (O) wells. (P) $T_n$-bottom means. (Q) Examples of $T_r$-bottom maps in empty wells. (R, S) Average radial profiles of $T_r$-bottom for small (R) and large (S) wells. (T) $T_r$-bottom means. (U) Examples of $T_c$-bottom maps in empty wells. (V, W) Average radial profiles of $T_c$-bottom for small (V) and large (W) wells. (X) $T_c$-bottom means. n=6 for each condition, from at least three independent experiments. Data are mean ± standard deviation. Scale bars: 5 µm.

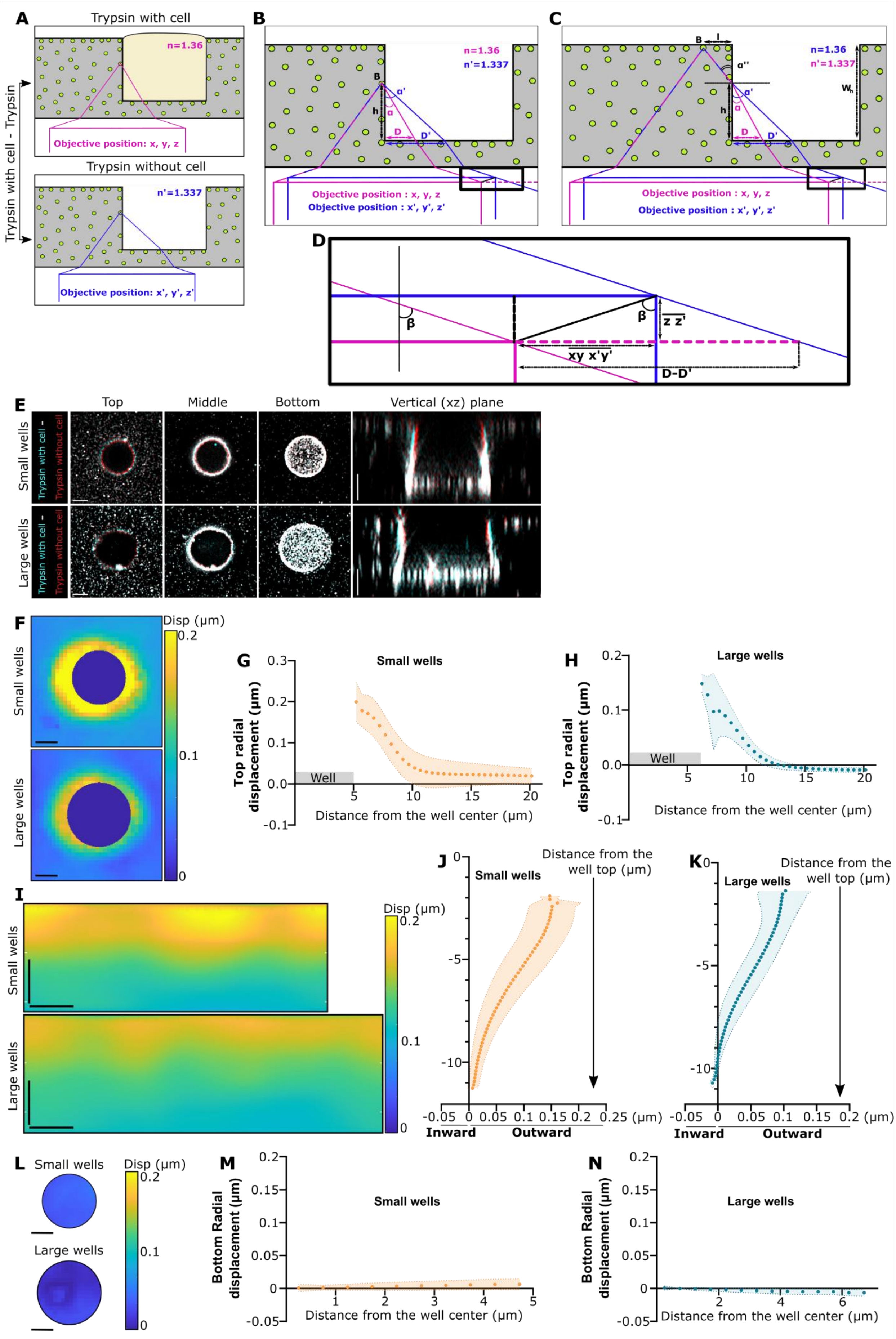

A
Trypsin with cell
n=1.36
Objective position: x, y, z
Trypsin without cell
n'=1.337
Objective position: x', y', z'
Trypsin with cell - Trypsin
B
n=1.36
n'=1.337
Objective position : x, y, z
Objective position : x', y', z'
C
n=1.36
n'=1.337
Objective position : x, y, z
Objective position : x', y', z'
D
β
z z'
xy x'y'
D-D'
E
Top
Middle
Bottom
Vertical (xz) plane
Small wells
Large wells
Trypsin with cell -
Trypsin without cell
F
Disp (µm)
Small wells
Large wells
G
Small wells
Top radial displacement (µm)
Well
Distance from the well center (µm)
H
Large wells
Top radial displacement (µm)
Well
Distance from the well center (µm)
I
Disp (µm)
Small wells
Large wells
J
Small wells
Distance from the well top (µm)
Inward
Outward
K
Large wells
Distance from the well top (µm)
Inward
Outward
L
Small wells
Large wells
Disp (µm)
M
Small wells
Bottom Radial displacement (µm)
Distance from the well center (µm)
N
Large wells
Bottom Radial displacement (µm)
Distance from the well center (µm)

**Figure S2: Characterization of the "refractive" field of displacement induced by the cell lensing effect in 15 kPa gels.** (A) Schematic of the experimental set-up. The lensing effect is measured by comparing two trypsinization conditions, the first one with the cell still present in the well, and the reference without it. (B) The laser beam used to image the beads is refracted differently in these conditions, as shown by the distinct laser paths drawn in pink (with cell) and blue (without cell). Consequently, the objective is displaced from the position (x, y, z) to the position (x', y', z') when imaging beads on the wall of the well. (C) same as (B) when imaging beads in the surface surrounding the well. (D) Zoom in on the displacement between the objective positions (x, y, z) and (x', y', z') present in panels (B) and (C). The theoretical calculation of the displacement is presented in Supplementary Information - note 2. (E) Superposition of the bead localization before (cyan) and after (red) cell removal from the well. Three xy planes of the stacks are shown (Top, Middle, and Bottom of the well) as well as a vertical plane (xz) of the same stacks. (F) Examples of radial displacement maps of the top surface. (G, H) Average radial profile of displacements on the top surface for small (G), and large (H) wells. (I) Example maps of normal wall displacement. (J, K) Average profiles of normal wall displacement along the z-axis for the small (J), and large (K) wells. (L) Example maps of bottom displacements. (M, N) Average radial profiles of bottom displacements for small (M) and large (N) wells. For (G, H, J, K, M, N) data are mean ± standard deviation. n=9 and 10 wells for small and large wells respectively, from at least three independent experiments. Scale bars: 5 µm.

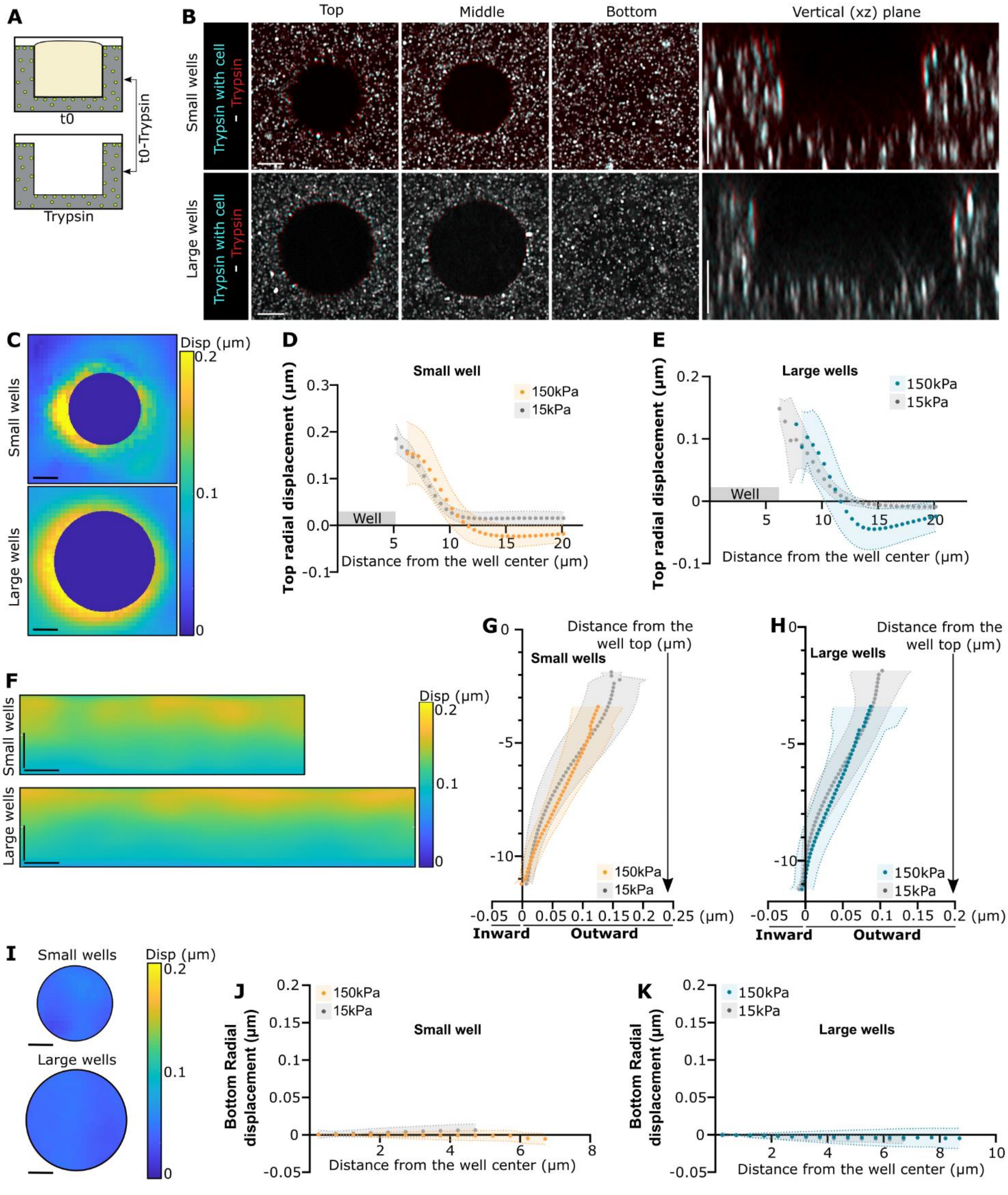


**Figure S3: Characterization of the "refractive" field of displacement induced by the cell lensing effect in 150 kPa gels.** (A) Schematic of the experimental set-up. (B) Superposition of the bead localization before (cyan) and after (red) cell removal from the well. Three xy planes of the stacks are shown (Top, Middle, and Bottom of the well) as well as a vertical plane (xz) of the same stacks. (C) Example radial displacement maps of the top surface. (D, E) Average radial profiles of displacements on the top surface for small (D), and large (E) wells. (F) Example maps of normal wall displacement. (G, H) Average profiles of normal wall displacement along the z-axis for the small (G), and large (H) wells. (I) Example maps of bottom displacements. (J, K) Radial profiles of bottom displacements for small (J) and large (K) wells. In (D, E, G, H, J, K) 15 kPa lensing effect data from

Figure S2, Supplementary Information are shown in gray for comparison. Well dimensions in 15 and 150 kPa gels do not match due to differential swelling. For (D, E, G, H, J, K) data are mean ± standard deviation. n=22 and n=17 cells for small and large wells respectively, from at least three independent experiments. Scale bars: 5 µm.

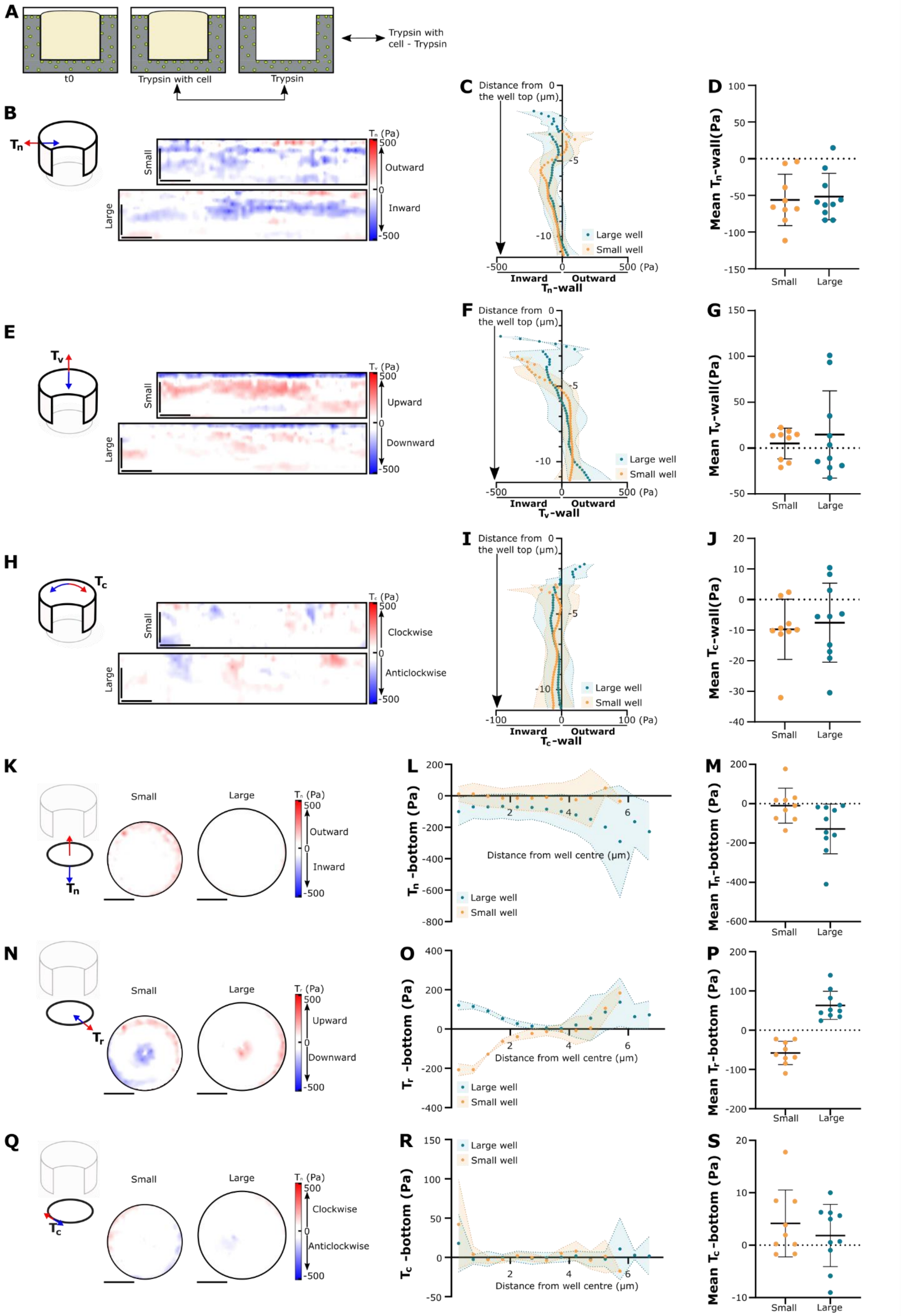

A
t0
Trypsin with cell
Trypsin
Trypsin with cell - Trypsin
B
$T_n$
Small
Large
$T_n$ (Pa)
500
Outward
0
Inward
-500
C
Distance from the well top (μm)
0
-5
-10
Large well
Small well
-500
0
500 (Pa)
Inward
Outward
$T_n$-wall
D
Mean $T_n$-wall(Pa)
100
50
0
-50
-100
-150
Small
Large
E
$T_v$
Small
Large
$T_v$ (Pa)
500
Upward
0
Downward
-500
F
Distance from the well top (μm)
0
-5
-10
Large well
Small well
-500
0
500 (Pa)
Inward
Outward
$T_v$-wall
G
Mean $T_v$-wall(Pa)
150
100
50
0
-50
Small
Large
H
$T_c$
Small
Large
$T_c$ (Pa)
500
Clockwise
0
Anticlockwise
-500
I
Distance from the well top (μm)
0
-5
-10
Large well
Small well
-100
0
100 (Pa)
Inward
Outward
$T_c$-wall
J
Mean $T_c$-wall(Pa)
20
10
0
-10
-20
-30
-40
Small
Large
K
$T_n$
Small
Large
$T_n$ (Pa)
500
Outward
0
Inward
-500
L
$T_n$ -bottom (Pa)
200
0
-200
-400
-600
-800
2
4
6
Distance from well centre (μm)
Large well
Small well
M
Mean $T_n$-bottom (Pa)
200
0
-200
-400
-600
Small
Large
N
$T_r$
Small
Large
$T_r$ (Pa)
500
Upward
0
Downward
-500
O
$T_r$ -bottom (Pa)
400
200
0
-200
-400
2
4
6
Distance from well centre (μm)
Large well
Small well
P
Mean $T_r$-bottom (Pa)
200
100
0
-100
-200
Small
Large
Q
$T_c$
Small
Large
$T_c$ (Pa)
500
Clockwise
0
Anticlockwise
-500
R
$T_c$ -bottom (Pa)
150
100
50
0
-50
2
4
6
Distance from well centre (μm)
Large well
Small well
S
Mean $T_c$-bottom (Pa)
20
10
0
-10
Small
Large

**Figure S4: Measurement of the traction noise in experiments corrected for the cell lensing effect.** (A) Schematic of the experimental set-up: trypsinized cells still within wells (trypsin with cell) were compared to wells after removing the cells (trypsin without cell). (B) Examples of $T_n$-wall maps. (C) Average $T_n$-wall profiles along the z-axis. (D) $T_n$-wall mean. (E) Examples of $T_v$-wall maps. (F) Average $T_v$-wall profile along the z-axis. (G) $T_v$-wall mean. (H) Examples of $T_c$-wall maps. (I) Average $T_c$-wall profile along the z-axis. (J) $T_c$-wall mean. (K) Examples of $T_n$-bottom maps. (L) Average Radial profiles of the $T_n$-bottom. (M) $T_n$-bottom mean. (N) Examples of $T_r$-bottom maps. (O) Average Radial profiles of the $T_r$-bottom. (P) $T_r$-bottom mean. (Q) Examples of $T_c$-bottom maps. (R) Average Radial profiles of the $T_c$-bottom. (S) $T_c$-bottom mean. Data are shown as mean ± standard deviation. n=9 and 10 cells for small and large wells respectively, from at least three independent experiments. Scale bars: 5 µm.

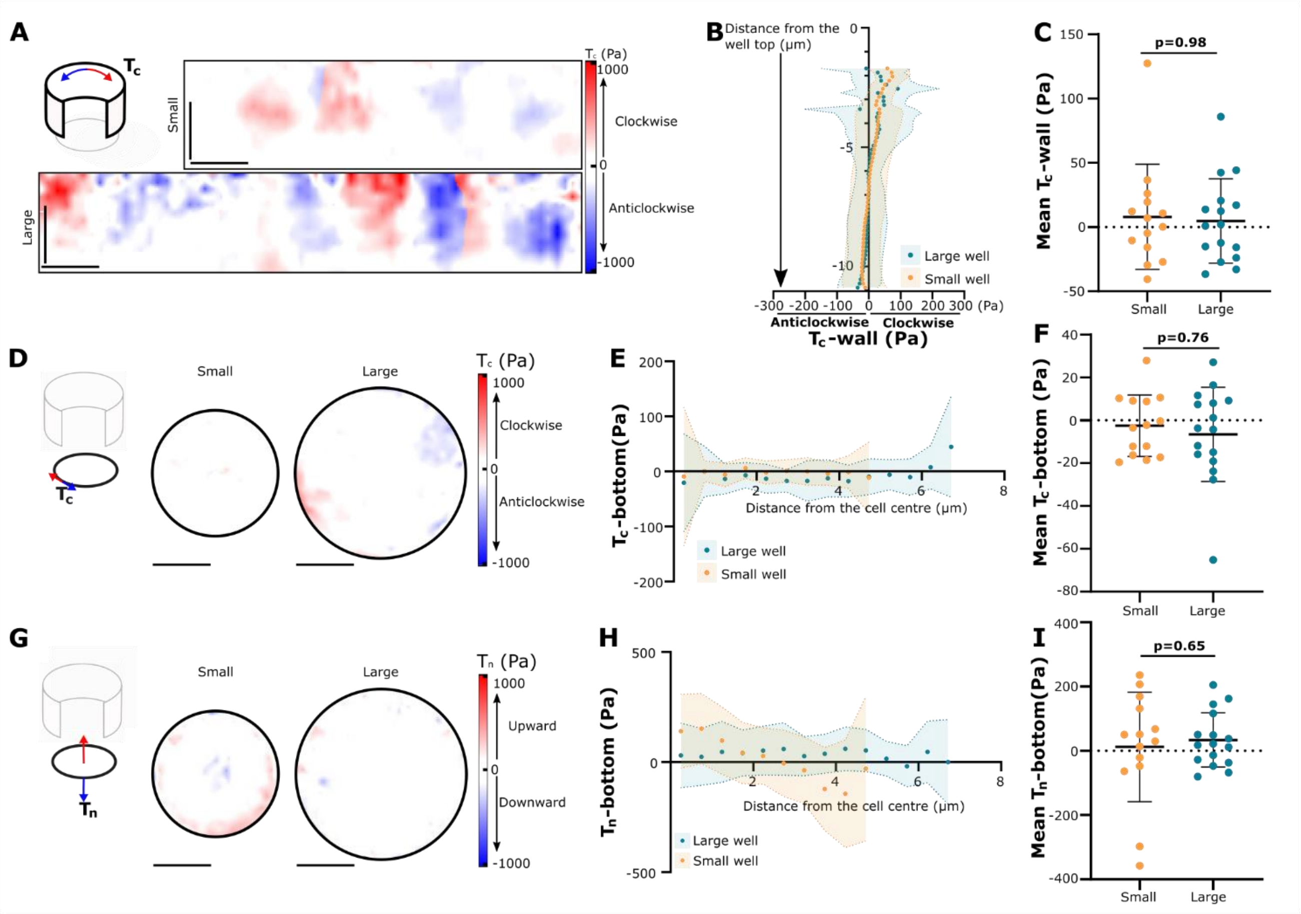


**Figure S5: Wall circumferential tractions and bottom normal and circumferential tractions for cells placed on small or large wells.** (A) Examples of $T_c$-wall maps. (B) Average $T_c$-wall profile along the z-axis. (C) $T_c$-wall mean. p=0.98, Mann-Whitney test. (D) Example $T_c$-bottom maps. (E) Average radial profiles of the $T_c$-bottom. (F) $T_c$-bottom mean. p=0.76, Mann-Whitney test. (G) Example $T_n$-bottom maps. (H) Average radial profiles of the $T_n$-bottom. (I) $T_n$-bottom mean. p=0.65, unpaired t-test. Data are shown as mean ± standard deviation. n=14 and n=16 for small and large wells respectively, from at least three independent experiments. Scale bars: 5 μm.

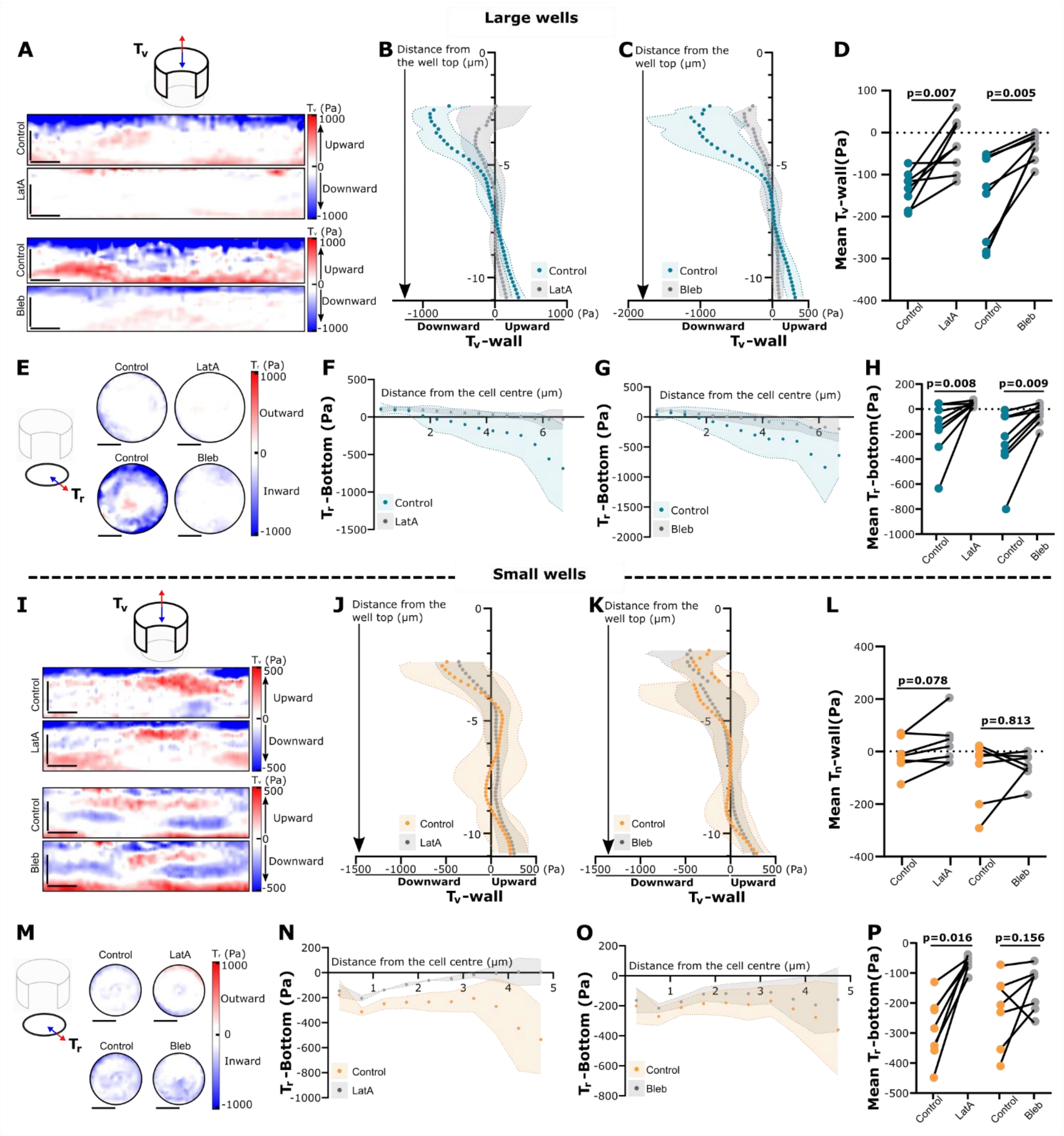


**Figure S6: Wall vertical tractions and bottom radial tractions for cells placed on small or large wells.** (A, I) Example $T_v$-wall maps comparing the blebbistatin/latrunculin A condition to the control for large (A), and small (I) wells. (B, C, J, K) Average $T_v$-wall profiles along the z axis for large (B, C), and small (J, K) wells. Data are mean ± standard deviation. (D, L) $T_v$-wall means for large (D), and small (L) wells. Mean paired data are represented as connected. For large wells, Control/LatA: p=0.008, paired t-test; Control/Bleb: p=0.005, paired t-test. For small wells, Control/ LatA: p=0.078, Wilcoxon matched-pairs signed rank test; Control/Bleb: p=0.813, Wilcoxon matched-pairs signed rank test. (E, M) Example $T_r$-bottom maps comparing the blebbistatin/latrunculin A condition to the control for large (E), and small (M) wells. (F, G, N, O) Average $T_r$-bottom profiles along the z-axis for large (F, G), and small (N, O) wells. Data are mean ± standard deviation. (H, P) $T_r$-bottom means for large (H), and small (P) wells. Mean paired data are represented as connected. For large wells, Control/LatA: p=0.008, Wilcoxon matched-pairs signed rank test; Control/Bleb: p=0.009, paired t-test. For small wells, Control/ LatA: p=0.016, Wilcoxon matched-pairs signed rank test; Control/Bleb: p=0.156,

Wilcoxon matched-pairs signed rank test. n=8 for each large well condition, n=7 for each small well condition, from at least three independent experiments. Scale bars: 5 µm.

### Finite Element method for traction inference

Laura M. Faure[1,†,*], Manuel Gómez-González[1,†], Ona Baguer[1], Jordi Comelles[1,3], Elena Martínez[1,2,3], Marino Arroyo[1,4,5,6], Xavier Trepat[1,2,7,8,*], Pere Roca-Cusachs[1,7,*]

*[1]Institute for Bioengineering of Catalonia (IBEC), Barcelona Institute of Science and Technology (BIST), Barcelona, Spain*
*[2]Centro de Investigación Biomédica en Red en Bioingeniería, Biomateriales y Nanomedicina (CIBER-BBN), Barcelona, Spain*
*[3]Department of Electronics and Biomedical Engineering, University of Barcelona (UB), Barcelona, Spain*
*[4]Laboratori de Càlcul Numèric (LaCàN), Universitat Politècnica de Catalunya (UPC), Barcelona, Spain*
*[5]Institut de Matemàtiques de la UPC–BarcelonaTech (IMTech), Barcelona, Spain*
*[6]Centre Internacional de Mètodes Numèrics en Enginyeria (CIMNE), Barcelona, Spain*
*[7]University of Barcelona (UB), Barcelona, Spain*
*[8]Institució Catalana de Recerca i Estudis Avançats (ICREA), Barcelona, Spain*

#### Direct approach

The problem of traction inference from displacement data (TFM) can be implemented in different ways depending on the context [1]. Here, given the geometric complexity of wells, and possibly the geometric and material nonlinearity of the gel, we resort to the finite element method (FEM) for mechanical calculations. Given measurements of the displacement field on the surface of the gel, the most direct approach is to impose it as a boundary condition in the free surface of a 3D finite element model of the gel [2, 3]. The imposition of surface displacements generates tractions in the gel, which are readily available in the FEM and which correspond to the sought after tractions exerted by cells on the gel. This direct approach is compatible with nonlinear elasticity [4].

In our setting, we call $S_{\text{top}}$ the top surface of the gel and $S_{\text{well}}$ the well surface, including the lateral and bottom surfaces, Figure 1. The gel volume is denoted by $V$. We denote the surface displacement field measured on these two parts of the gel surface by $\boldsymbol{u}^*_{\text{top}}$ and $\boldsymbol{u}^*_{\text{well}}$.

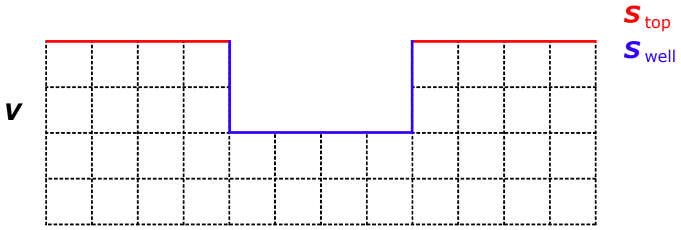


**Figure 1:** Simplified representation of the computational domain: bulk of the gel ($V$, black), top surface of the gel ($S_{\text{top}}$, red) and well surface ($S_{\text{well}}$, blue).

*Corresponding author
†Equal authorship.

The direct approach mentioned above can be formalized as follows. Consider the mechanical boundary value problem of finding the displacement field $\boldsymbol{u}(x, y, z)$ defined over $V$ such that

$$\nabla \cdot \boldsymbol{\sigma} = \boldsymbol{0} \quad \text{in} \quad V, \tag{1}$$

$$\boldsymbol{u} = \boldsymbol{u}^*_{\text{top}} \quad \text{on} \quad S_{\text{top}}, \tag{2}$$

$$\boldsymbol{u} = \boldsymbol{u}^*_{\text{well}} \quad \text{on} \quad S_{\text{well}}, \tag{3}$$

where the first equation states force balance in the interior of the gel, and $\boldsymbol{\sigma}$ is the Cauchy stress tensor, which follows from the elastic constitutive equation as a function of the gradient of the displacement field. We also impose boundary conditions in the bottom (zero displacements) and lateral sides (traction-free) of the computational domain. The FEM discretizes these equations to obtain a system of equations whose unknowns are the displacements at the nodes of the finite element mesh. In the case of linear elasticity, this system of equations is linear, wheres for nonlinear elasticity, it is necessary to solve a nonlinear systems of equations iteratively. Once $\boldsymbol{u}$ has been obtained in $V$, the stress tensor can be computed and the surface tractions exerted on the gel obtained as [1]

$$\boldsymbol{T}_{\text{top}} = \boldsymbol{\sigma} \cdot \boldsymbol{n} \quad \text{on} \quad S_{\text{top}}, \tag{4}$$

$$\boldsymbol{T}_{\text{well}} = \boldsymbol{\sigma} \cdot \boldsymbol{n} \quad \text{on} \quad S_{\text{well}}, \tag{5}$$

where $\boldsymbol{n}$ is the outer normal field to the surface.

We implemented this approach in Abaqus (Dassault Systemes), considering a Neo-Hookean nonlinear hyperelastic model. We defined the geometry as a square prism with the same hight than the gel and a cylindrical cavity, on the top surface, of the same depth and average radius than the physical well. The edges of the cylinder were rounded using a radius of 0.5 µm for the bottom edge and of 1 µm for the top edge of the cylinder to closely follow the geometry of the physical wells and avoid artefactual accumulation of traction at sharp edges. Each model was meshed in Abaqus with hybrid 4-node linear tetrahedron elements. An illustrative mesh and map of inferred surface tractions is shown in Figure 2a.

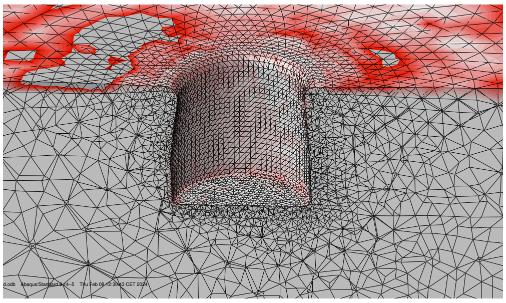

**(a)** Representative mesh and traction field calculated with the Direct approach.

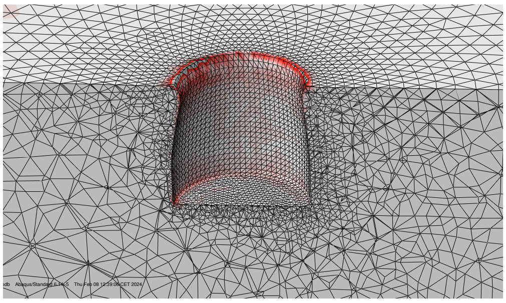

**(b)** Representative mesh and traction field calculated with the Inverse approach.

**Figure 2:** Representative traction fields calculated with the Direct and Inverse approaches.

The tractions resulting from this approach have an obvious problem. Because fibronectin only coats the well, cells are only able to adhere and exert tractions on $S_{\text{well}}$. However, the computational method also predicts tractions on the top surface $S_{\text{top}}$. There is thus an obvious violation of force balance on $S_{\text{top}}$. A simple way to address this is to consider a modified mechanical problem with imposed displacements on $S_{\text{well}}$ and traction-free boundary conditions on $S_{\text{top}}$,

$$\nabla \cdot \boldsymbol{\sigma} = 0 \quad \text{in} \quad V, \tag{6}$$

$$\boldsymbol{\sigma} \cdot \boldsymbol{n} = \boldsymbol{0} \quad \text{on} \quad S_{\text{top}}, \tag{7}$$

$$\boldsymbol{u} = \boldsymbol{u}^*_{\text{well}} \quad \text{on} \quad S_{\text{well}}, \tag{8}$$

and compute the traction on the well surface as in Eq. (5). Although this alternative direct approach localizes cell tractions where they should be, it does not use the measurements on the top surface $\boldsymbol{u}^*_{\text{top}}$.

##### Inverse approach

We develop next an inverse approach [5] to infer the surface traction that uses all the experimentally available information and whose resulting solution satisfies mechanical equilibrium in the bulk and at the surfaces. To formalize this method, we assume isotropic linear elasticity, but it is readily generalizable to anisotropic or nonlinear elasticity [6]. In the linear case, the strain tensor $\boldsymbol{\varepsilon}$ can be computed in terms of the displacement field and the stress tensor in terms of the strain tensor as

$$\varepsilon_{ij} = \frac{1}{2}\left(\frac{\partial u_i}{\partial x_j} + \frac{\partial u_j}{\partial x_i}\right) \quad \text{and} \quad \sigma_{ij} = \lambda \varepsilon_{kk}\delta_{ij} + 2\mu\varepsilon_{ij}, \tag{9}$$

where $\lambda$ and $\mu$ are the Lamé elastic coefficients.

In the inverse method, the unknowns are the traction field $\boldsymbol{T}_{\text{well}}$ on the well surface $S_{\text{well}}$ and the displacement field $\boldsymbol{u}$ in the gel. The goal is to minimize the discrepancy between $\boldsymbol{u}$ and the measured displacements, quantified by the quadratic penalty functional

$$\mathcal{E}(\boldsymbol{u}) = \frac{1}{2}\int_{S_{\text{top}}} ||\boldsymbol{u} - \boldsymbol{u}^*_{\text{top}}||^2 dS + \frac{1}{2}\int_{S_{\text{well}}} ||\boldsymbol{u} - \boldsymbol{u}^*_{\text{well}}||^2 dS. \tag{10}$$

The displacement field and the surface tractions should be consistent with mechanical equilibrium, which is imposed as constraints using Lagrange multipliers, resulting in the following Lagrangian functional

$$\begin{aligned} \mathcal{L}(\boldsymbol{u}, \boldsymbol{T}_{\text{well}}; \boldsymbol{\lambda}, \boldsymbol{\mu}_{\text{top}}, \boldsymbol{\mu}_{\text{well}}) = &\frac{1}{2}\int_{S_{\text{top}}} ||\boldsymbol{u} - \boldsymbol{u}^*_{\text{top}}||^2 dS + \frac{1}{2}\int_{S_{\text{well}}} ||\boldsymbol{u} - \boldsymbol{u}^*_{\text{well}}||^2 dS \\ &+ \int_V \boldsymbol{\lambda} \cdot (\nabla \cdot \boldsymbol{\sigma}(\boldsymbol{u}))\, dV \\ &+ \int_{S_{\text{top}}} \boldsymbol{\mu}_{\text{top}} \cdot \boldsymbol{\sigma}(\boldsymbol{u}) \cdot \boldsymbol{n}\, dS + \int_{S_{\text{well}}} \boldsymbol{\mu}_{\text{well}} \cdot (\boldsymbol{\sigma}(\boldsymbol{u}) \cdot \boldsymbol{n} - \boldsymbol{T}_{\text{well}})\, dS \\ &+ \frac{\alpha}{2}\int_{S_{\text{well}}} ||\boldsymbol{T}_{\text{well}}||^2 dS + \frac{\beta}{2}\int_{S_{\text{well}}} ||\nabla \boldsymbol{T}_{\text{well}}||^2 dS. \end{aligned} \tag{11}$$

The stress tensor $\boldsymbol{\sigma}(\boldsymbol{u})$ is an explicit function of the displacements according to Eq. (9). The integral in the second line imposes mechanical equilibrium within the gel, the first integral in the third line imposes mechanical equilibrium on the top surface (traction-free boundary condition), and the second integral imposes mechanical equilibrium between the unknown surface tractions and the stresses in the material on the well surface, with $\boldsymbol{\lambda}$, $\boldsymbol{\mu}_{\text{well}}$ and $\boldsymbol{\mu}_{\text{well}}$ being the corresponding Lagrange multiplier fields. The integrals in the fourth line are regularization functionals penalizing the magnitude and the variations of the tractions at the well with regularization parameters $\alpha$ and $\beta$. The equations allowing us to compute the unknowns and the Lagrange multipliers are obtained by making $\mathcal{L}(\boldsymbol{u}, \boldsymbol{T}_{\text{well}}; \boldsymbol{\lambda}, \boldsymbol{\mu}_{\text{top}}, \boldsymbol{\mu}_{\text{well}})$ stationary with respect to all of its arguments.

Regularization is know to be required in such inverse methods [5]. If regularization parameters are too small or zero, then the solution is dominated by noise. If they are too large, then regularization functionals bias the optimization procedure by trying to make tractions too small or too uniform, inducing a large discrepancy between computational and measured displacement (large $\mathcal{E}(\boldsymbol{u})$). Optimal parameters can be estimated by plotting $\mathcal{E}(\boldsymbol{u})$ and the regularization functionals as a function of the regularization parameters, and selecting parameters that achieve a tradeoff between these conflicting objectives.

Physically, the inverse approach is more reasonable because it imposes exactly the balance laws (here mechanical equilibrium), which should apply independently of the material behavior and the accuracy of the measurements, and instead, it allows a small discrepancy between the inferred displacements and the measured ones, which are subjected to experimental error. In contrast, the direct approach imposes the noisy measurements exactly and satisfies force balance at the top surface only approximately.

To implement the inverse method in the linear case, we exported the stiffness matrix corresponding to the finite element mesh $\mathbf{K}$ from Abaqus into Matlab to form a discrete finite element version of the continuous Lagrangian in Eq. (11). Given the array of nodal displacements $\mathbf{u}$, the matrix-vector product $\mathbf{Ku}$ returns the internal forces at the nodes resulting from the material elasticity. Mechanical equilibrium at each node and along the $x$, $y$ or $z$ direction (corresponding to a degree of freedom labelled by $I$) depends on the balance of $\sum_{J\in D} K_{IJ}u_J$ and an externally applied force $f_I$, which in our case are only present in the well. Here, $D$ denotes the set of all degrees of freedom of the finite element model. Therefore the equations $\sum_{J\in D} K_{IJ}u_J = 0$ hold for all degrees of freedom $I$ except those corresponding to a node in the well surface, and are the discrete version of Eqs. (6,7). Defining the set of degrees of freedom corresponding to the top surface as $D_{\text{top}}$ and those corresponding to the well surface as $D_{\text{well}}$, the discrete Lagrangian can be reformulated as

$$\begin{aligned} L(\mathbf{u}, \mathbf{f}; \boldsymbol{\lambda}, \boldsymbol{\mu}) = & \frac{1}{2} \sum_{I\in D_{\text{top}}} \left(u_I - u^*_{\text{top},I}\right)^2 + \frac{1}{2} \sum_{I\in D_{\text{well}}} \left(u_I - u^*_{\text{well},I}\right)^2 \\ & + \sum_{I\in D} \lambda_I \left(\sum_{J\in D} K_{IJ}u_J - f_I\right) + \sum_{I\in\notin D_{\text{well}}} \mu_I f_I \\ & + \frac{\alpha}{2} \sum_{I\in D_{\text{well}}} f_I^2 + \frac{\beta}{2} \sum_{I\in D_{\text{well}}} (\nabla f)_I^2 . \end{aligned} \tag{12}$$

The first sum in the second line constrains the external nodal forces $f_I$ to agree with the internal forces and the second sum constrains the external forces to be non-zero only at the surface of the well. We note that all terms in the sums can be multiplied by appropriate integration weights to make the correspondence with the continuous Lagrangian more quantitative, but it is not required and this form is clearer. The last term in Eq. (12) controls the magnitude of the surface gradients of the traction force, computed using the finite element basis functions of the triangulation corresponding to the surface of the well.

The stationarity conditions

$$\partial L/\partial u_I = 0 \quad \text{for} \quad I \in D \tag{13}$$

$$\partial L/\partial f_I = 0 \quad \text{for} \quad I \in D \tag{14}$$

$$\partial L/\partial \lambda_I = 0 \quad \text{for} \quad I \in D \tag{15}$$

$$\partial L/\partial \mu_I = 0 \quad \text{for} \quad I \notin D_{\text{well}} \tag{16}$$

result in a linear system of equations, whose solution provides all unknowns including the external forces $f_I$ at the surface of the well.

As previously discussed, the regularization parameters should achieve a tradeoff between regularity of the inferred tractions and error in the measured displacements. We found good results considering only the gradient regularization, and therefore we ignored the first regularization term ($\alpha = 0$) to minimize bias. To select $\beta$, we sampled a range of values and, for each choice, we computed the error function

$$E(\mathbf{u}) = \frac{1}{2} \sum_{I \in D_{\text{top}}} \left(u_I - u^*_{\text{top},I}\right)^2 + \frac{1}{2} \sum_{I \in D_{\text{well}}} \left(u_I - u^*_{\text{well},I}\right)^2, \tag{17}$$

and the gradient regularization function

$$G(\mathbf{f}) = \frac{1}{2} \sum_{I \in D_{\text{well}}} (\nabla F)^2_I \,. \tag{18}$$

As $\beta$ increases, the discrepancy between measured and computational displacements measured by the error function in Eq. (17) is expected to grow, whereas the gradient regularization function in Eq. (18) decreases since lager $\beta$ biases the solution towards smoother tractions at the expense of fitting the data. Our computational results confirm these expectations, Fig. 3. These kind of figure allows us to select an appropriate value of $\beta$, achieving a tradeoff between the conflicting objectives.

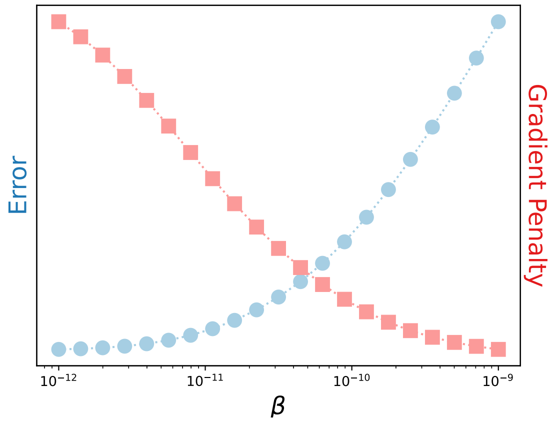


**Figure 3:** Error of the solution and gradient penalty as a function of $\beta$, for a representative well.

To convert nodal forces into tractions $f_I$, we divide them by the surface area corresponding to that particular node, Figure 4. For each node at the surface, we consider all the element faces at the surface to which it belongs. For each face, we calculate the area fraction dividing its area by the number of nodes in it. For the example in Figure 4, the area fraction would correspond to one third of the face area. The surface corresponding to a node would then be the sum of all of the area fractions of the faces to which it belongs.

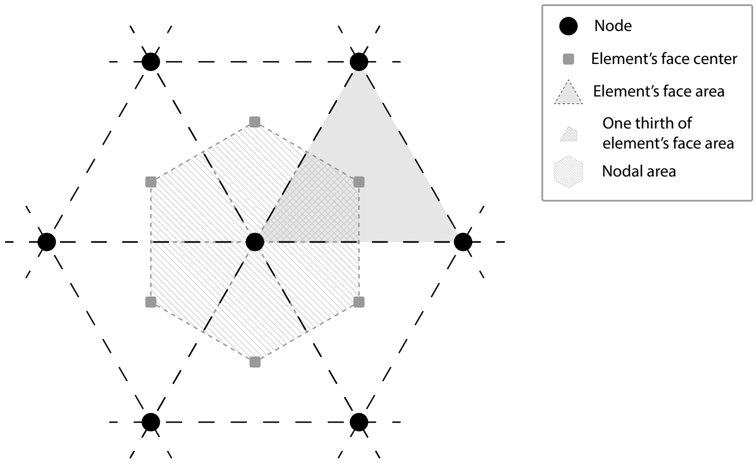


**Figure 4:** Representation of the surface area corresponding to each node.

**Supplementary information - note 2: Theoretical and experimental estimation of lensing effects and their correction.**

Our experimental setup contains materials with different refractive indexes. This includes the culture medium (n’ =1,337)[1], polyacrylamide gels ($n_{gel}$ =1,349)[2], the immersion medium of the objective (glycerin, $n_{gly}$=1,456, Zeiss Immersol – Ref: 462959-9901-000) and MCF10A cells, for which previous measurements showed n=1.36[3]. Thus, as cells do not have the same refraction index as their surrounding environment they act as lenses, modifying the light path. This “lensing effect” modifies the apparent location of the fluorescent markers in the gels, even for cells that are not applying any forces, for example, trypsinized cells. Given the regular shape of our wells, this effect can be both predicted theoretically and measured experimentally.

*Theoretical calculation*

We will estimate here the apparent displacement between the images (that is, positions of the objective) of a fluorescent bead (B) imaged successively through the cell (position x, y, z, light path displayed in pink) or culture media (position x’,y’,z’ light path displayed in blue) (Figure S2B, D, Supplementary Information). To compare with our experiments, we will use the characteristics of our 40X glycerin immersion objective, with a numerical aperture (NA) of 1.2.

We first consider the case of beads located along the wall of the well. The following equations can be written (see Figure S2B-D, Supplementary Information for the definitions of angles α, α’, α’’ and β and distances h, D, D’, $\overline{xy\,x'y'}$, $\overline{z\,z'}$):

(1) $NA = n_{gly}\sin(\beta) = n sin(\alpha) = n'\sin(\alpha') = n_{gel}\ \sin(\alpha'')$

(2) $h = \frac{D}{\tan(\alpha)} = \frac{D\prime}{\tan(\alpha')}$

(3) $\overline{xy\,x'y'} = \frac{D-D\prime}{2}$

(4) $\overline{z\,z'} = \frac{\overline{xy\,x\prime y\prime}}{\tan(\beta)}$,

Combining equations (1), (2), (3), and (4) we get:

(5) $\overline{xy\,x'y'} = \frac{h}{2}\left(\frac{\frac{NA}{n}}{\sqrt{1-\frac{NA^2}{n^2}}} - \frac{\frac{NA}{n'}}{\sqrt{1-\frac{NA^2}{n'^2}}}\right)$

(6) $\overline{z\,z'} = \frac{h}{2}\frac{\left(\frac{\frac{NA}{n}}{\sqrt{1-\frac{NA^2}{n^2}}} - \frac{\frac{NA}{n'}}{\sqrt{1-\frac{NA^2}{n'^2}}}\right)}{\frac{\frac{NA}{n_{gly}}}{\sqrt{1-\frac{NA^2}{n_{gly}^2}}}}$

Equations (5) and (6) give respectively the apparent horizontal displacement ( $\overline{xy\,x'y'}$) and vertical displacement ($\overline{z\,z'}$), both proportional to the vertical distance between the bead and the bottom of the well $h$. This means that we will observe larger displacement when we image closer to the surface of the gel. Using our known values of NA and n', and the previously reported value of n for MCF10A cells (n=1.36), and for $h = 11\ \mu m$, we estimate $\overline{xy\,x'y'} = 0.88\ \mu m$ and $\overline{z\,z'} = 0.34\ \mu m$. Of note, we expect such displacements to be measurable along the xy but not z direction, where the resolution of our microscope is lower.

For beads located along the surface of the gel, we consider that:

$$(7)\ \frac{l}{\mathrm{W}_h - \mathrm{h}} = \tan(\alpha'')$$

Where l is the horizontal distance from the well edge, $W_h$ is the total height of the well, and α'' is the angle of the refracted light beam in the gel (Figure S2C, Supplementary Information). Combining this equation with (1), (5), and (6), we get:

$$(8)\ \overline{xy\,x'y'} = \left(\frac{W_h}{2} - l\frac{*n\sqrt{1-\frac{NA^2}{n^2}}}{2n_{gel}\sqrt{1-(\frac{n^2}{n_{gel}^2}(1-\frac{NA^2}{n^2}))}}\right) * \ (\frac{\frac{NA}{n}}{\sqrt{1-\frac{NA^2}{n^2}}} - \frac{\frac{NA}{n'}}{\sqrt{1-\frac{NA^2}{n'^2}}})$$

$$(9)\ \overline{zz'} = \left(\frac{W_h}{2} - l\frac{n\sqrt{1-\frac{NA^2}{n^2}}}{2n_{gel}\sqrt{1-(\frac{n^2}{n_{gel}^2}(1-\frac{NA^2}{n^2}))}}\right)\frac{(\frac{\frac{NA}{n}}{\sqrt{1-\frac{NA^2}{n^2}}} - \frac{\frac{NA}{n'}}{\sqrt{1-\frac{NA^2}{n'^2}}})}{\frac{\frac{NA}{n_{gly}}}{\sqrt{1-\frac{NA^2}{n_{gly}^2}}}}$$

Again, we get a linear relationship between l and the horizontal and vertical displacements. The displacements are maximum at the well edge, and then decrease until reaching zero at:

$l = \frac{W_h * n_{gel}\sqrt{1-(\frac{n^2}{n_{gel}^2} * \left(1-\frac{NA^2}{n^2}\right))}}{n\sqrt{1-\frac{NA^2}{n^2}}}$, approximately 20 μm from the well edge. For a total well height of 11μm, displacements at the well edge are the same mentioned above.

*Experimental measurements*

Following the theoretical predictions, we tested if we could experimentally measure the displacement induced by the lensing effect, we focused on the xy displacements as we did not detect displacements along the z-axis, as expected. To do so, we disrupted adhesions between the cell and the gel through trypsinization, thereby eliminating traction forces while keeping the cell in the well for imaging. We then compared the image stack of the fluorescent beads to that obtained after flushing the cell out of the well (Figure S2A, E, Supplementary Information). We performed this experiment in small and large wells and obtained 3D displacement vectors for each voxel of our stacks. We then analyzed the normal displacements to the wall of the well, the radial displacements in the top surface, and the radial displacements in the bottom of the well. Normal displacements to the wall of the well were negligible at the bottom of the well and increased with height, reaching values of the order of 0,1-0,14 μm at the top (Figure S2E, I-K, Supplementary Information). Displacements pointed away from the well centre. Radial displacements in the top surface showed a maximum displacement near the edge of the well (0,1-0,2 μm), followed by a decay toward 0 at 5 μm from the well edge (Figure S2F-H, Supplementary Information). Radial displacements at the bottom of wells were negligible (Figure S2L-N, Supplementary Information). These displacement profiles were highly reproducible, depended on well size, and followed theoretical predictions, equations (5), (6), (8), and (9)): along the wall well, they increased in an approximately linear way from the bottom to the top of the wells, along the top surface, they decreased also linearly from the well edge, and on the bottom surface, they were negligible. Interestingly, maximum values for the normal displacements to the wall were between 0.1 and 0.14 μm, which is significantly below the 0.88 μm prediction at a height of 10 μm. Similarly, the radial top displacement became negligible around 5 μm from the well edge compared to the 20 μm predicted. This suggests that, in our setting, the refraction indexes of cells, medium, and gel are closer to each other than previously reported[3].

Importantly, we confirmed these experimental trends by measuring the apparent displacements generated by cells when placed on very stiff gels, here 150 kPa, that cannot be deformed by cells (Figure S3, Supplementary Information). The consistency between both approaches (measurements in trypsinized cells on 15 kPa gels, versus non-trypsinized cells on 150 kPa gels) confirmed that the results were due to the lensing effect, and not potential residual tractions in trypsinized cells.

*Correction of lensing effect*

To correct for the lensing effect, we approximated it by a characteristic "refractive" field of displacement. This was taken as the mean observed displacements caused by lensing, as measured in Figure S2, Supplementary Information. Of note, we used 15 kPa (Figure S2, Supplementary Information) rather than 150 kPa (Figure S3, Supplementary Information) data for this, as 150 kPa gels had a slightly different geometry than 15 kPa gels (the stiffness used throughout the manuscript) due to different swelling. We considered the "refractive" displacements to be additive to any displacement caused by mechanical forces exerted by cells on the substrate. Thus, we subtracted the "refractive" displacement field from displacements measured in our system before computing tractions (Figure 1B).

Finally, we evaluated the noise given by this correction. We measured the tractions of trypsinized cells. As these cells exert no forces on the wells, the only bead displacement they generate should be associated with the lensing effect (Figure S4A, Supplementary Information). After the lensing effect correction, the result corresponds to the remaining noise and gives us information on the variability between each individual case and the average value. As in empty wells, the highest peaks of traction are located at the well edge while the rest of the maps did not present any characteristic pattern (Figure S4, Supplementary Information). These peaks were higher than the one observed for the empty wells, reaching magnitude of 500 Pa instead of the 100 Pa seen for instance in the $T_v$-wall of the empty wells (Figure S4E-G, Supplementary Information). The correction of the lensing effect is maximal at the top of the well, hence, this is the position where we expect the most noise between the individual wells and the mean value (Figure S4B, C, E, F, H, I, Supplementary Information). Thus, the correction of the lensing effect, which was maximal at the top of the well, adds an extra noise to the one given by gel swelling. The maps of traction at the well bottom were also characterized by some accumulation of forces at the edge, particularly for the $T_n$-bottom of the large wells (Figure S4K, N, Q, Supplementary Information). This could be explained by the fact that we have a lower resolution measuring displacement along the z-axis than in x,y.
